# A co-proteomic view of metabolite-specific interactions in the *Botrytis cinerea-Arabidopsis* pathosystem

**DOI:** 10.64898/2026.06.05.730517

**Authors:** Anna Jo Muhich, Celine Caseys, Brooke Grabbe, Christian Montes-Serey, Justin Walley, Daniel J. Kliebenstein

**Affiliations:** Department of Plant Sciences, University of California Davis, Davis CA 95616, USA; Department of Plant Pathology and Microbiology, Iowa State University, Ames, IA 50011, USA

## Abstract

To successfully infect their myriad hosts, generalist plant pathogens must tolerate a vast arsenal of plant specialized defense metabolites. To understand how host-specific metabolites influence plant-generalist pathogen interactions, we conducted a co-proteomic analysis of both *Arabidopsis thaliana* and *Botrytis cinerea* proteomes from the same samples during early infection. The Arabidopsis proteomic responses to Botrytis center around induction and suppression of defense metabolite pathways, particularly camalexin and glucosinolates. Several Botrytis proteins involved in key virulence pathways were induced within 32-48 hours, including potential defense metabolite detoxification proteins. Co-proteomic analysis using a panel of Arabidopsis genotypes with differing glucosinolate profiles revealed that disruptions to the glucosinolate pathway had broad changes on the Arabidopsis proteome, and that Botrytis induces specific proteins in response to presence/absence of Arabidopsis defense metabolites. Among the proteins that were induced quickly on infection and linked to the presence of glucosinolates, we validated a novel isothiocyanate hydrolase in Botrytis, BcSaxA, that catabolizes isothiocyanates in vitro. Gene expression data further indicated BcSaxA is expressed only in dicot hosts containing isothiocyanates. Our study describes a highly dynamic host proteome during infection with Botrytis and elucidates metabolite-specific infection strategies for a generalist pathogen.

## Introduction

In host-generalist systems, pathogen virulence and host defense is often highly polygenic and many factors determine disease outcomes. Host plants defend against attacking pathogens with a blend of conserved resistance pathways and lineage-specific defenses (Sucher et al. 2020; Singh et al. 2026a). Similarly, generalist pathogens use both wide ranging strategies and specific responses to each host to coordinate a successful infection (Kusch et al. 2022; Singh et al. 2026b). This generates a pathosystem where both the host and pathogen utilize numerous mechanisms, including small RNAs, proteins, and toxic metabolites. Of these interactive components, a particularly intriguing aspect is the plant hosts’ use of lineage-specific defense metabolites, and the pathogen’s response. Diversity of these specialized metabolites across land plants is vast, with over 200,000 unique structures reported spanning a growing number of classifications (Osbourn and Lanzotti 2009). While generalist pathogens are frequently assumed to rely on broad-spectrum resistance mechanisms, recent work is showing that they contain tolerance mechanisms that are deployed in a host-specific manner (Westrick et al. 2021). This raises the potential that generalist pathogens induce specific tolerance mechanisms to handle the hosts’ metabolic diversity.

*Botrytis cinerea*, hereafter Botrytis, is a model generalist necrotrophic pathogen that is beginning to be found to contain specific defenses against host specialized metabolites. A first category of detoxification mechanisms uses efflux transporters to remove potentially toxic metabolites from pathogen cell space. These often have broad substrate binding capacity, for example, the ATP-binding cassette (ABC) transporter BcatrB conducts efflux of multiple structurally unrelated host compounds including resveratrol, camalexin, and eugenol (Schoonbeek et al. 2001; Schoonbeek et al. 2003; Stefanato et al. 2009), and is upregulated in response to structurally diverse compounds spanning Solanaceae, Fabaceae, and Brassicaceae (Bulasag et al. 2023). Additionally, Major Facilitator Superfamily (MFS) transporters have been shown to provide tolerance for multiple host compounds, including BcmfsG in efflux of isothiocyanates (ITCs) in Arabidopsis (Vela-Corcía et al. 2019), and Bcmfs1 for the alkaloid camptothecin and the perylenequinone cercosporin (Hayashi et al. 2002). A second category involves detoxification of host metabolites through enzymatic degradation. Contrasting with efflux transport, this strategy may be more compound class specific. For example, this has been demonstrated in Botrytis in response to resveratrol (Schouten et al. 2002), tomatine (Quidde et al. 1998; You et al. 2024), and rishitin (Bulasag et al. 2024). The short chain dehydrogenase, BcCPDH, has been shown to degrade capsidiol in pepper (B. Ward 1972), but is not induced when tested against similarly structured host metabolites (Kuroyanagi et al. 2022). The growing body of literature showing a blend of both general and highly specialized Botrytis responses to host metabolites highlights the need for understanding how changes in host metabolism shape host-Botrytis interactions as a whole.

However, plant-pathogen interactions are highly dynamic, and examining either organism in isolation can obscure how these responses are interconnected. The pathogen is not merely a bystander responding with detoxification mechanisms. It also actively suppresses host defense metabolism, leading to transcriptional downregulation of pathways involved in defense-associated metabolite production (Cai et al. 2023; Singh et al. 2023). Simultaneous omics approaches that profile both host plant and pathogen within the same sample are therefore critical for understanding these interactive processes. Co-transcriptomic analysis across diverse Botrytis isolates infecting 10 eudicot hosts demonstrated that, alongside a conserved set of genes with similar expression across the hosts, Botrytis also has sets of genes with host-specific expression (Singh et al. 2026b), indicating a large degree of transcriptional programs conditional on the host. Similarly, host responses rely heavily on lineage-specific transcriptional reprogramming, with only a limited core of conserved defenses (Singh et al. 2026a). In the Arabidopsis-Botrytis pathosystem, co-transcriptomics enabled identification of links between host defense pathways and fungal virulence modules (Zhang et al. 2019). While transcriptomic studies have provided substantial insight, they are inherently limited in their ability to capture protein abundance and function due to post-transcriptional regulation. Although recent advances have expanded proteomic analyses of Botrytis (Coste et al. 2025), the use of co-proteomics to simultaneously characterize both host plant and pathogen proteomes remain relatively unexplored and have thus far been applied primarily in studies with Fusarium (Fabre et al. 2019; Fabre et al. 2021; Witzel et al. 2024)

In this study, we conduct a co-proteomic analysis of Botrytis cinerea model isolate B05.10 with Arabidopsis to better understand how lineage specific defense metabolites are altered by and alter host-pathogen interactions, using the Brassicales-specific metabolites glucosinolates (GSLs) as a model. We combined an early infection time course and multiple Arabidopsis genotypes with differing GSL profiles to understand how these defense metabolites shape host-generalist interactions. We show that, like transcription, Arabidopsis proteomic responses to Botrytis center around induction and suppression of specific metabolic pathways. We further demonstrate Botrytis induces specific proteins in response to presence/absence of Arabidopsis defense metabolites. Among these specifically upregulated proteins, we identify a novel ITC hydrolase in Botrytis, BcSaxA, and demonstrate it breaks down Arabidopsis ITCs in vitro. Our results contribute to a growing understanding of the role of host metabolite specificity in plant-generalist pathogen interactions.

## Results

### Arabidopsis-Botrytis co-proteomics time course

To characterize temporal patterns of Arabidopsis-Botrytis interaction during early infection and identify a suitable timepoint to test for defense metabolite mutant effects on the interaction, we conducted a Arabidopsis-Botrytis co-proteomics time course assay, generating global proteomes from mock- and infected Col-0 leaf tissue at 8-hour intervals from 0-48 HPI. The raw dataset identified 11,712 Arabidopsis proteins and 1,548 Botrytis proteins. After filtering for stochastic detection and low abundance proteins, 11,669 Arabidopsis and 1,537 Botrytis proteins remained (Supplementary Table 1). Principal component analysis (PCA) showed that Arabidopsis proteomes are clustered by timepoint and infection status. At 48 HPI, infected Arabidopsis proteomes began to be clearly separated from mock samples along PC1, which explained 66.6% of the variance (Supplementary Figure 1A). The mock samples allowed for identification of a minimal baseline level of detected proteins associated with the predicted Botrytis proteome. These appear to be from either conserved proteins shared across fungi or a residual level of Botrytis (Supplemental Figure 1B and Supplementary Table Raw Data). Given the absence of any visible infections in any mock samples, we presume this to be a residual signal. Using this mock baseline, there began to be a signal where infected samples diverged along PC1 (97.5% of the variance) starting at 16 HPI with stronger signals starting at 32 HPI (Supplementary Figure 1B).

### Arabidopsis coordinates proteomic shifts during early infection with Botrytis

To investigate the temporal patterns in Arabidopsis protein abundance during Botrytis infection, we modeled each protein with the formula *Protein ∼ Infected + Time + Infected x Time*. This allowed for comparison of infection response to mock samples at each timepoint to adjust for any diurnal variation or responses to the assay that may exist (Choudhary et al. 2015; Krahmer et al. 2022). A total of 10,542 proteins were modeled with this approach, of which 2,355 showed significant effects of infection or the Infection x Time interaction after multiple testing correction (p < 0.05; Supplementary Table 2). Hierarchical clustering of these infection-responsive proteins identified 20 clusters with different temporal abundance patterns (Figure 1 and 2). Several clusters displayed pronounced infection responses. Cluster 11 showed the strongest upregulation and was enriched for proteins involved in camalexin biosynthesis and JA response (Figure 2). Other upregulated clusters were largely associated with indole defense metabolism (cluster 10 for tryptophan and cluster 2 for indole glucosinolates) or JA signaling (Cluster 7). This suggests that proteome induction is largely focused on the production of defense metabolites. The most strongly downregulated cluster, cluster 14, was enriched for proteins associated with aliphatic GSL biosynthesis, potentially to facilitate increases in indolic defense metabolites. The other largely downregulated cluster, cluster 9, was involved in cell wall metabolism. These patterns are consistent with prior transcriptomic time course analysis of Arabidopsis infected with Botrytis (Windram et al. 2012), although transcriptional responses were reported to occur earlier, with camalexin biosynthesis induced by 14 HPI and abiotic stress responses declining by 24 HPI. Most of the remaining clusters exhibited modest differences between mock and infected samples, often with peak divergence at 48 HPI. Overall, Arabidopsis proteins respond to Botrytis in a highly coordinated manner during early infection, with a strong emphasis on fluctuation of specialized metabolic pathways.

**Figure 1.**
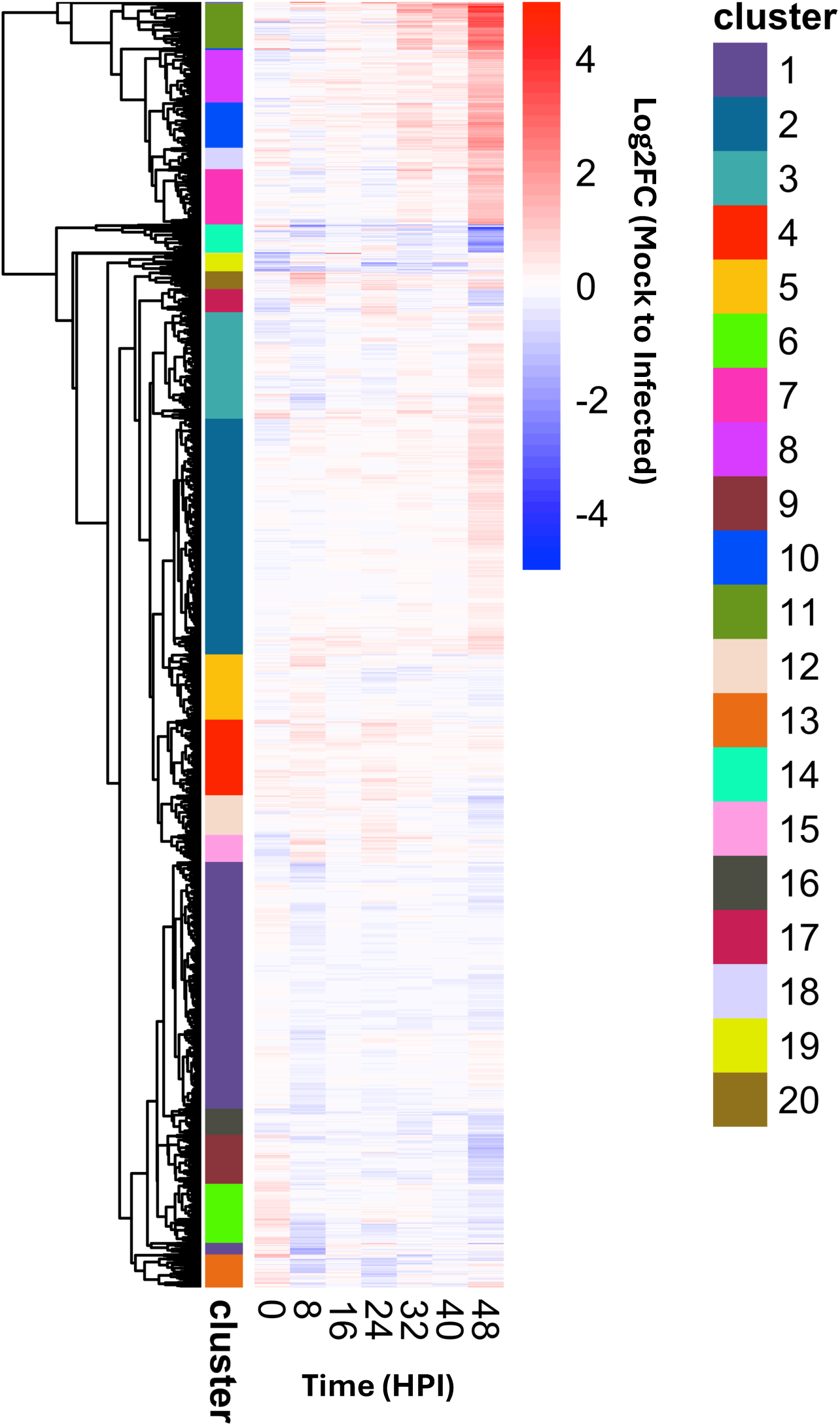
Heatmap of Arabidopsis protein abundance changes over time (log2FC) for proteins significantly affected by Botrytis infection or the infection × time (HPI) interaction (p < 0.05). Hierarchical clustering was performed using complete linkage, and clusters were defined with dynamic tree cutting. A total of 27 outlier proteins with extreme abundance patterns were removed prior to clustering. For visualization, extreme log2FC values were capped at ±5.

**Figure 2.**
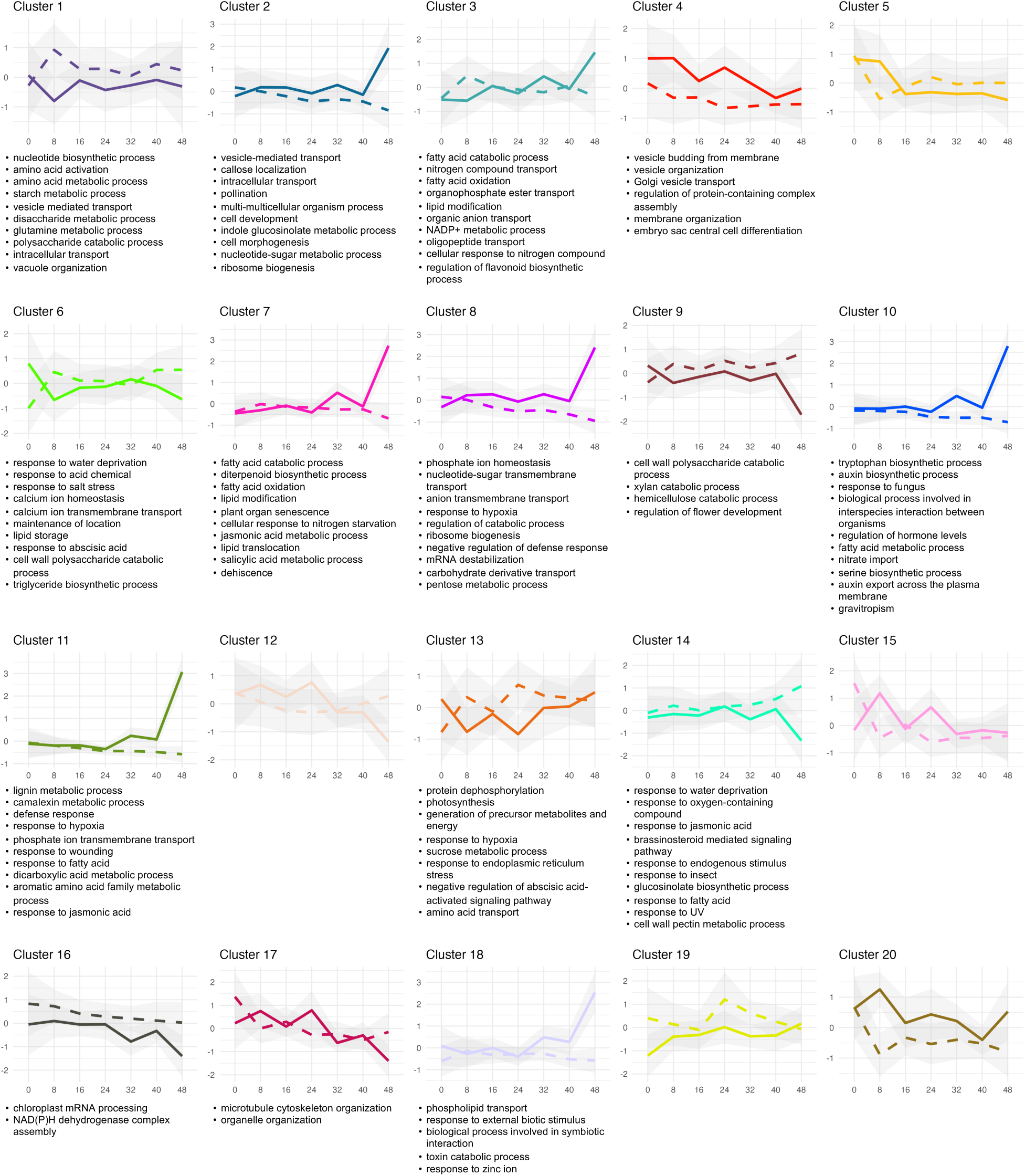
Arabidopsis mean protein abundances from clusters (generated in Figure 1) across time course infection with Botrytis isolate B05.10. Solid lines represent mean abundances for infected tissue, while dashed lines represent mean abundances of mock. Standard deviations are shown as shadows. Significantly enriched (p < 0.05) GO terms within each cluster are listed. Clusters with no terms listed had no significant enrichment.

The proteomics along with previous metabolite and transcriptomic analysis all indicate that Arabidopsis has a focused response to Botrytis using defense metabolism. Specifically, indolic defense metabolism is induced while the aliphatic glucosinolate metabolism is decreased. To focus on these changes, we conducted a pathway level analysis of camalexin, indolic glucosinolate, aliphatic glucosinolate biosynthesis and catabolism proteins. A small subset of 27 proteins with outlying abundance patterns were excluded from the previous clustering, including key camalexin biosynthetic enzymes PAD3, CYP71A12, and CYP71A13. For more finescale understanding of these enzymes and temporal abundance of GSL biosynthetic proteins more broadly during Botrytis infection, known proteins from these pathways were plotted for abundance over time, regardless of whether they responded significantly to infection (Supplementary Figure 2). In agreement with the strong induction of camalexin accumulation, all camalexin biosynthetic proteins (PAD3, CYP71A12, and CYP71A13) were strongly upregulated starting between 24 and 32 HPI. Botrytis infection also leads to an induction of indolic glucosinolates which is associated with the upregulation of the key pathway specific proteins CYP79B3, CYP81F1, IGMT4, and SOT16 with about an 8-hour delay in comparison to camalexin biosynthetic protein induction. Apparently supporting this induction was the increase in proteins supplying sulfur (APSKinase) or core pathway proteins not previously ascribed as being indole specific (CSLyase, GSTU13 and UGT74B).

Aliphatic glucosinolate accumulation is reduced in front of a growing Botrytis lesion and infection lowers expression of the pathway (Kliebenstein et al. 2005; Zhang et al. 2015). Supporting this was the observed downregulation of key aliphatic GSL biosynthetic proteins MAM1, CYP79F, CYP83A. However, most other proteins in the aliphatic GSL pathway had relatively stable levels in response to infection (BCAT3/4, GSTF11, SOT17, SOT18, etc.) indicating that the decrease does not require shifts in the whole pathway. Investigating catabolism showed that PEN2 is highly upregulated. PEN2 is an atypical myrosinase that primarily catabolizes indole GSLs through an active process in living plant cells (Bednarek et al. 2009), differentiating it from the general myrosinases TGG1/2 that catabolize both indolic and aliphatic GSLs and were stable during infection. Interestingly, the catabolic protein classes with the most changes were the downstream specifier enzymes including nitrilases (NIT) and nitrile specifier proteins (NSP) suggesting a shift towards other downstream catabolites. Overall, our results show that defense metabolite, transcript and protein changes agree in response to Botrytis infection.

### Botrytis proteomic dynamics during infection

Botrytis proteins were quantified and modeled across the same time-course samples from 8-48 HPI (Supplementary Table 3). Detectable Botrytis protein abundance increased markedly beginning at 32 HPI and began to plateau by 48 HPI (Figure 3A). There were Botrytis proteins statistically significant above the mock threshold starting at 8 HPI with a gradual increase up to the most Botrytis proteins being first consistently detected at 40 HPI (Figure 3B). Proteins detected in the early timepoints (8, 16 HPI) were largely confined to those involved in fundamental cellular processes, dominated by core metabolic and ribosomal proteins (Supplementary Table 4). The rate of new protein identification significantly dropped at the 48 HPI time point suggesting that the Botrytis proteome is starting to saturate (Figure 3B). GO enrichment analysis of the proteins based upon when they were first induced only detected a significant enrichment at 16 and 24 hours and these were limited to broad, high-level biological process terms (Supplementary Figure 3), suggesting that early timepoints are largely detecting high-abundance general process proteins like the ribosome and lack the detection threshold to map fine-scale temporal changes in Botrytis protein abundance. At 48 HPI protein detection reached its maximum across the dataset (Figure 3C). To provide insight on known protein dynamics, we focused on the abundance of known Botrytis virulence factors (Figure 3D). This included biosynthetic enzymes in the botcinic acid pathway (BOA), the botrydial cluster (BOT), enzymatic cell death inducing proteins (CDIPs) Bcpg1, Bcpg2, and Bcxyn11A, and non-enzymatic CDIPs Bcnep1/2 and Bcspl1. PG1 but not PG2 was detected. Among the non-enzymatic CDIPs, only Bcspl1 was detected. Most Botrytis virulence proteins were first detected at 32 HPI with a general increase to 48 HPI. However, some variation was seen in the BOA proteins. Bcboa9, a polyketide synthase required for botcinic acid production (Dalmais et al. 2011), was detected at all timepoints, while others were not reliably detected until 48 HPI. Together, these results indicate 48 HPI shows detectable levels of major fungal virulence pathway proteins and is a suitable timepoint for downstream analysis of Botrytis proteome variation across Arabidopsis defense metabolite profiles.

**Figure 3.**
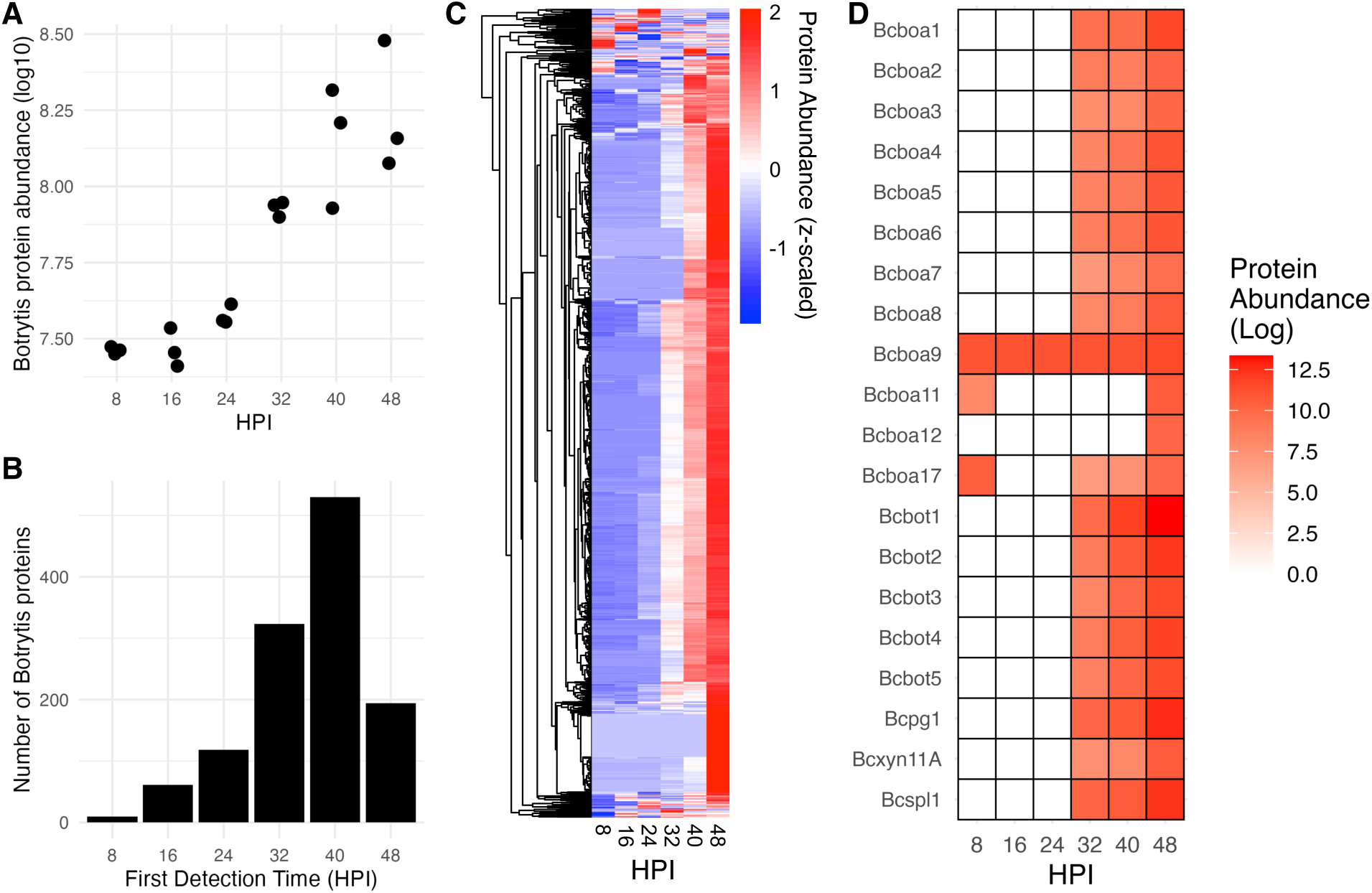
Botrytis proteome time course during infection of Arabidopsis Col-0. A) Raw Botrytis protein abundance collected from infected leaf tissue. B) Histogram of first detection time of each Botrytis protein. A protein was counted as detected if it appeared in two consecutive timepoints. C) Heatmap of total Botrytis protein abundance over time. Values were z-scaled within a protein. Dendrogram of proteins was generated by hierarchical clustering (average linkage) using Euclidean distance on protein abundances. D) Heatmap of protein abundance for selected Botrytis virulence factors over time. Non-zero values were log-transformed for visualization.

### Arabidopsis-Botrytis co-proteomics across host defense metabolite genotypes

Given the strong defense metabolite component in Arabidopsis responses to Botrytis and previous work suggesting generalist pathogens have metabolite-specific responses (Kuroyanagi et al. 2022; Kusch et al. 2022), we wanted to test how the plant’s defense metabolism influences the pathogen’s proteome. The above time-course analysis elucidated Arabidopsis-Botrytis proteomic shifts over time during infection and suggested that 48 HPI is a suitable timepoint for querying the Botrytis proteome and how it is altered by the plant’s defense metabolism. For the defense metabolite, we focused on the glucosinolates given the large collection of mutants available that alter this pathway. Co-proteomic profiling was performed at 48 HPI following infection of eight distinct Arabidopsis GSL genotypes with Botrytis isolate B05.10 (Table 1).

**Table 1.** Summary of samples collected for co-proteomics of Arabidopsis-Botrytis infection. Mock and infected tissue were collected for each above combination of genotype and timepoint and included 3 biological replicates.

| Arabidopsis Genotype | Mutant Background | Times Sampled |
| --- | --- | --- |
| Col-0 | WT | 0, 8, 16, 24, 32, 40, 48 HPI |
| Ler | WT | 48 HPI |
| Sha | WT | 48 HPI |
| *myb28* | *Col-0* | 48 HPI |
| *myb29* | *Col-0* | 48 HPI |
| *myb28/29* | *Col-0* | 48 HPI |
| *AOP2* | *Col-0* | 48 HPI |
| *tgg1/2* | *Col-0* | 48 HPI |

The panel included three well-studied wild-type accessions (Col-0, Ler, and Sha) alongside a series of mutants in the Col-0 background affecting the aliphatic GSL pathway (Figure 4A). First, these comprised the transcription factor mutants *myb28*, *myb29*, and *myb28/29* double mutant. Consistent with their regulatory roles, *myb28* has reduced short- and long-chained aliphatic GSLs, *myb29* has only reduced short-chained aliphatic GSLs, and the double mutant lacks aliphatic GSLs entirely (Sønderby et al. 2007).

**Figure 4.**
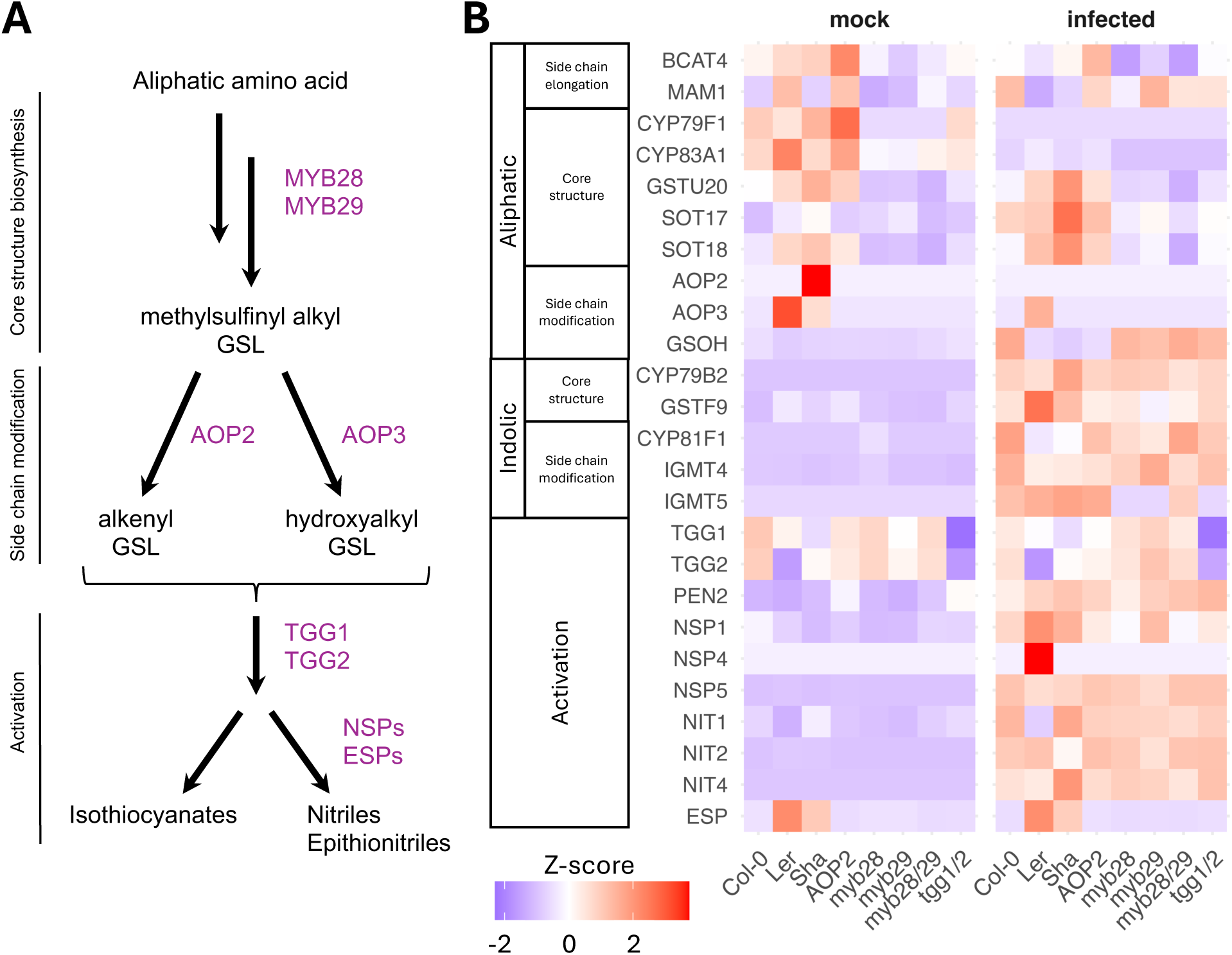
A) Simplified aliphatic glucosinolate pathway showing key roles of Arabidopsis mutant genotypes used in the study. B) Abundance of glucosinolate pathway enzymes across Arabidopsis accessions and glucosinolate mutant lines during infection with Botrytis isolate B05.10. Abundances were z-scaled within each protein. Some known pathway proteins were not detected in proteomic data and are not shown.

The three accessions provide diversity in in side-chain modification capacity mediated by AOP enzymes. Sha expresses AOP2, producing alkenyl GSL, while Ler expresses AOP3, producing hydroxylalkyl GSL. In contrast, Col-0 lacks functional AOP2 and AOP3 and accumulates methylsulfinylalkyl GSL precursors (Kliebenstein et al. 2001). We also included a “AOP2” complementation line that expresses a functional AOP2 in the Col-0 background from *Brassica oleracea* (Li and Quiros 2003; Wentzell et al. 2007).

Finally, we included the *tgg1/2* double mutant that abolishes the redundant TGG1 and TGG2 myrosinases that are predominantly responsible for catabolizing GSLs into isothiocyanates within the Col-0 background (Barth and Jander 2006). This mutant enables assessment of host and pathogen proteomic shifts in near absence of GSL breakdown.

Raw co-proteomic data from the UC Davis Proteomics Core Facility of these 8 Arabidopsis genotypes infected with Botrytis recovered 4,082 Arabidopsis proteins and 1,234 Botrytis proteins at 48 HPI. After stochastic detection and low abundance filtering as described above, 4,055 Arabidopsis proteins and 1,076 Botrytis proteins were suitable for analysis (Supplementary Table 5).

### Aliphatic GSL genotypes alter Arabidopsis proteome during infection

We initially tested expected proteomic differences among the Arabidopsis defense metabolite genotypes during Botrytis infection by investigating the abundances of key GSL pathway proteins (Figure 4B). As seen in the time course, early aliphatic GSL biosynthetic enzymes, including BCAT4, MAM1, CYP79F1, and CYP83A1, were generally downregulated following infection at 48 HPI in the wildtype accessions. As expected, the aliphatic GSL proteins were highly diminished in the double mutant (*myb28/29*) that does not accumulate these GSLs. Interestingly, the single *myb28* and *myb29* mutants also had measurably diminished protein accumulation even though they have at best a 50% decrease in metabolite (Gigolashvili et al. 2007; Sønderby et al. 2010). This lower baseline level attenuated any response to infection within the *myb28/29* mutants for the aliphatic GSL proteins. As expected for the side-chain modifying enzymes, AOP2 was expressed in Sha, AOP3 in Ler and GSOH in Col-0. This maps to their known presence/absence phenotypes for the causal genes in these accessions (Kliebenstein et al. 2001; Hansen et al. 2008). Interestingly, GSOH abundance increased during infection in Col-0 and the Col-0 background knockout mutants, but not the transgenic AOP2 line (also in the Col-0 background).

Consistent with time course analysis, indolic GSL pathway enzymes displayed stronger induction during infection than aliphatic pathway enzymes. This trend was broadly conserved across Arabidopsis genotypes, although the magnitude of induction varied most strongly in the wild-type accessions. Several activation enzymes, including PEN2, NSP1, NSP5, and NIT1/2/4, accumulated to higher levels during infection whereas others (TGG1/2, ESP) remained comparatively stable. Most of these responses were consistent across genotypes. However, PEN2 was more strongly upregulated in myb28/29 and tgg1/2 than Col-0, consistent with its role in indolic GSL hydrolysis.

We also analyzed the full proteomic data to model each defense metabolite genotype’s effect on all detected Arabidopsis proteins, and test if the mutants alter the response to infection with Botrytis. This allowed us to query how the plant’s defense metabolism may influence the infection response. Arabidopsis protein abundance was modeled relative to Col-0 for each comparison to sum the total number of proteins significantly different across genotype, infection, and the genotype x infection interaction (Supplementary Figure 4; Supplementary Table 6). Infection status was the main driver of proteomic variation, with approximately half of the detectable proteome responding to infection. However, in each comparison, hundreds of proteins still varied across genotype and the genotype’s interaction with infection. This genotype effect was predictably largest across the three wild-type accessions, followed by *myb28/29* knockouts with 1,377 proteins changing due to MYB28/29 status. While the AOP2 and *tgg1/2* comparisons showed considerably fewer proteins responding to genotype (849 and 722, respectively), this still indicates a large overall proportion of the proteome is affected by the presence of these GSL pathway enzymes.

For a more focused analysis of how the defense metabolite genotypes alter responses to infection, strongly infection-responsive proteins (>= 2 fold change, p < 0.05) in each GSL mutant comparison with wild type Col-0 were grouped (Figure 5). A large set of 411 infection-responsive proteins were shared between the mutants, representing a core conserved defense response to Botrytis unaffected by GSL pathway perturbation. However, each mutant displayed distinct proteomic signatures in response to infection. Functional enrichment of protein groups showed all three groups had alterations in other GSL biosynthetic proteins, but these proteins were distinct between AOP2 and myb28/29;tgg1/2. Further, myb28/29 and tgg1/2 both showed changes in upstream sulfur and indole metabolism. Overall, infection responses between mutants are largely shared and mainly differ in a few related processes between genotypes.

**Figure 5.**
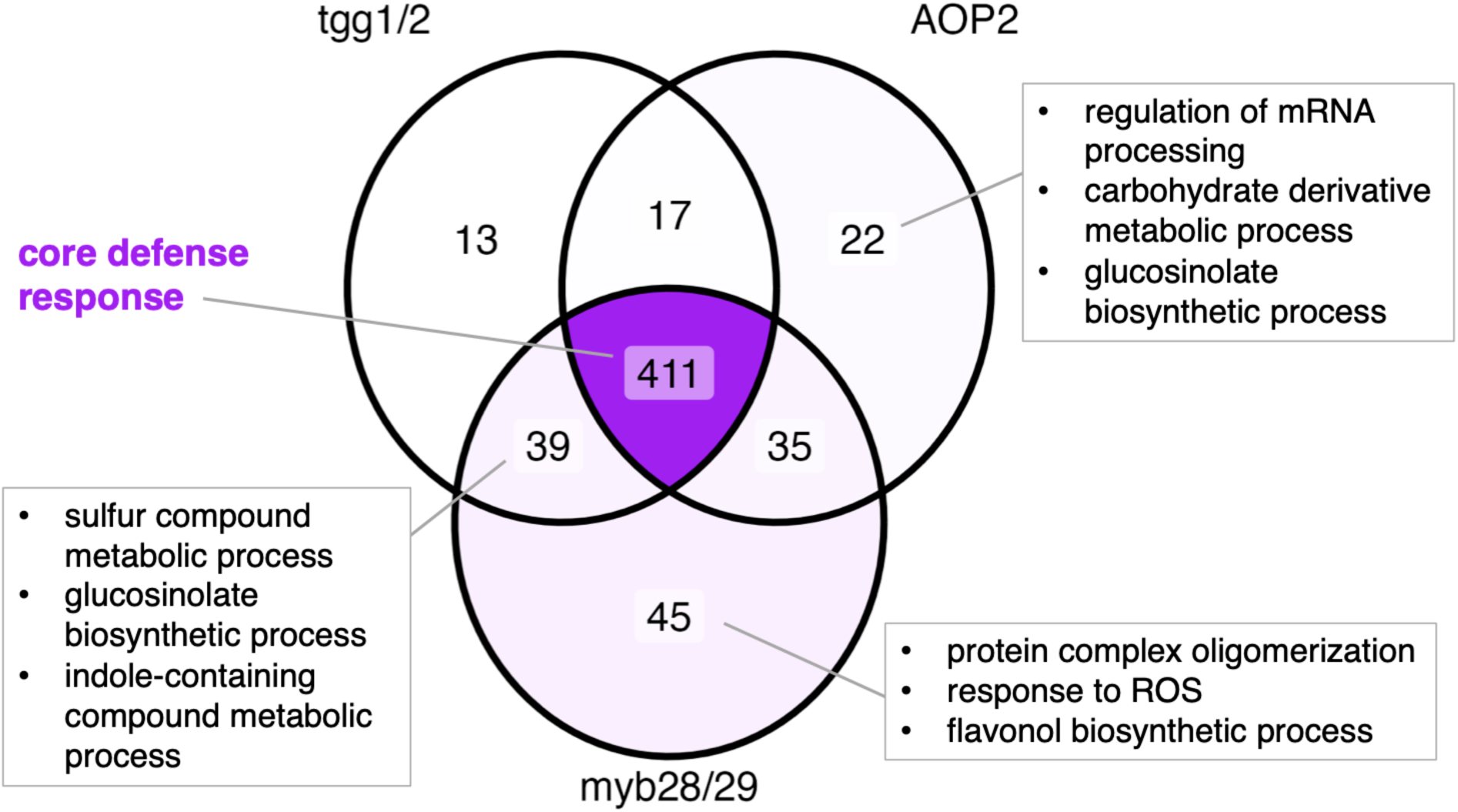
Arabidopsis proteins differentially accumulated across mutant comparisons. Venn diagram showing Arabidopsis proteins significantly responding to infection or genotype x infection (p < 0.05). Proteins are filtered for >= 2-fold change in any sample from Col-0 control (mock). Selected significant GO terms for each group are shown where applicable. The central core defense response denotes proteins that change in response to infection in all tested genotypes.

### Botrytis proteins show specialized responses to changes in aliphatic GSL profiles

To assess if Botrytis has proteomic responses specific to host metabolite variation, we analyzed the fungal proteomes during infection of the Arabidopsis genotypes. Protein abundance was modeled relative to an infection on Col-0 for each genotype comparison (Supplementary Table 7). To focus on Botrytis proteins with large genotype-dependent changes, a ≥ 2-fold change threshold was applied to filter through the statistically significant proteins. This identified 21 Botrytis proteins differentially abundant across the wild-type accessions, 13 across the *myb* mutants, 7 between AOP2 and Col-0, and 2 between *tgg1/2* and Col-0 (Figure 6).

**Figure 6.**
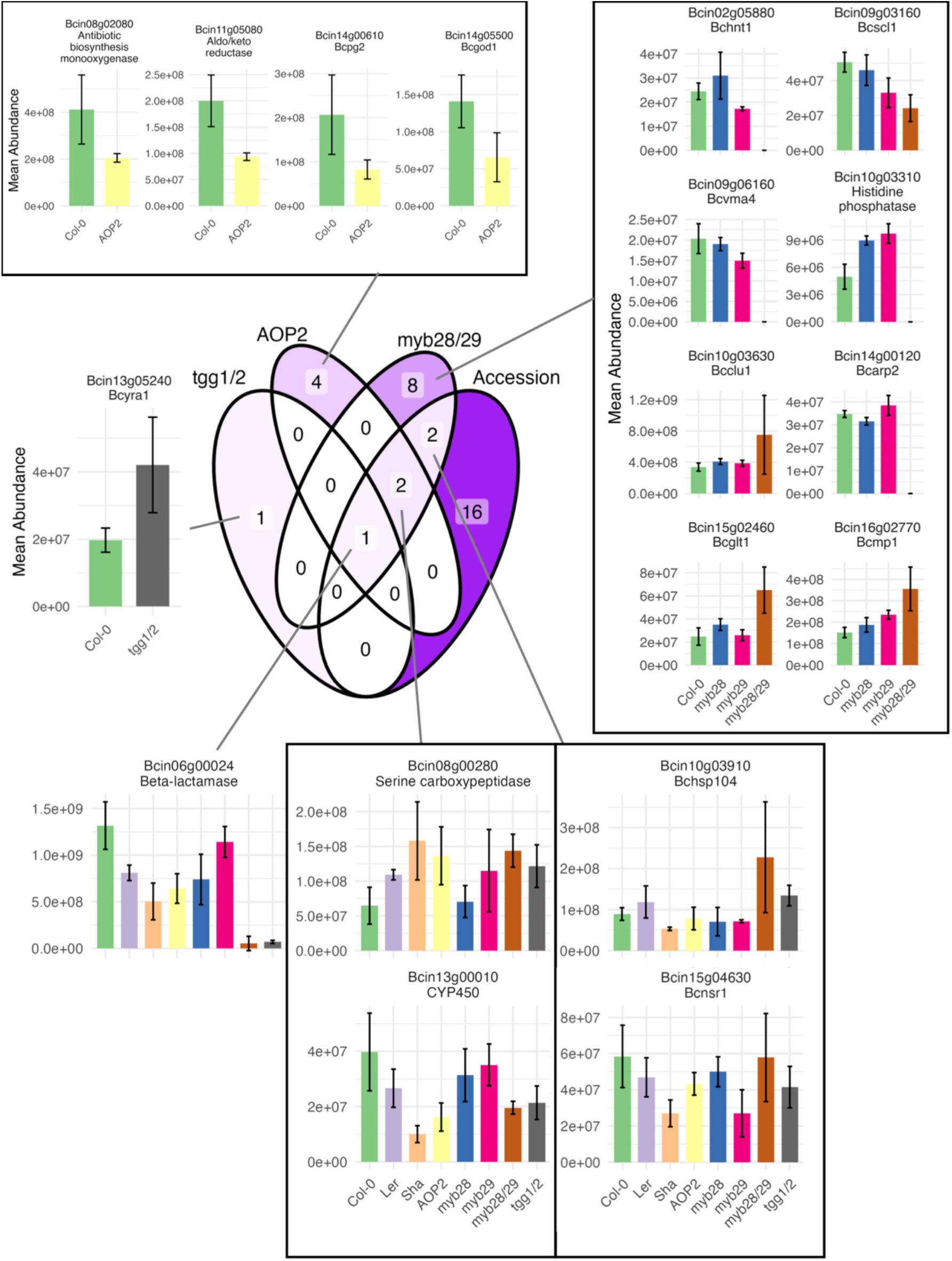
Venn diagram of Botrytis proteins with significantly altered accumulation during infection of different Arabidopsis accessions and glucosinolate mutants (p < 0.05; >= 2-fold change). Each model (protein abundance ∼ genotype) was performed independently for each comparison with Col-0. Mean and SD of protein abundances for each significant comparison are shown. The 16 proteins altered across accessions are shown in Figure S6.

Endopolygalacturonase Bcpg2 was significantly reduced in AOP2, along with glucose oxidase Bcgod1, an antibiotic biosynthesis monoxygenase Bcin08g02080, and an aldo/keto reductase Bcin11g05080. Bcpg2 is a reported host-specific virulence factor (Kars et al. 2005; Blanco-Ulate et al. 2014) that is upregulated in the presence of pectate as a carbon source (Zhang et al. 2014). Both Bcpg2 and the glucose oxidase Bcgod1 are also downregulated in the absence of transcription factor Bcxyr1, a key regulator of cell wall degrading enzymes (Ma et al. 2022). Although Bcxyr1 was not detected here, the reduction of Bcpg2 suggests that its accumulation may also respond to host defense metabolite profiles, potentially contributing to its host specificity.

In the *myb28/28* mutant, which lacks aliphatic GSLs, several fungal proteins were either absent or increased in abundance. Four Botrytis proteins were fully absent when infecting the *myb28/29* mutant: HIT-domain containing Bchnt1, ATPase subunit protein Bcvma4, histidine phosphatase Bcin10g03310, and actin binding protein Bcarp2. Conversely, four proteins were upregulated: clustered mitochondria family protein Bcclu1, glutamate synthase Bcglt1, metalloprotease Bcmp1, and ATPase Bchsp104. These proteins span diverse functional categories, suggesting that aliphatic GSLs may broadly influence Botrytis physiology.

Sixteen Botrytis proteins varied exclusively among the three WT Arabidopsis accessions. Notably, the efflux transporter BcatrB, known to export camalexin and other phenolic compounds (Schoonbeek et al. 2001; Schoonbeek et al. 2003; Stefanato et al. 2009), was reduced in Ler and Sha compared to Col-0 (Supplementary Figure 5). While camalexin accumulation differs across accessions during infection (Denby et al. 2004), its induction is highly time-dependent, and BcatrB may also respond to additional host-derived compounds.

Of these, there was only one protein, a metallo-ꞵ-lactamase (Bcin06g00024), that differed significantly across all genotype comparisons (Figure 6). It did not accumulate in plants lacking aliphatic GSLs and their isothiocyanate breakdown products, highlighting it as an interesting candidate tolerance mechanism for GSL-derived toxins. The closest homolog to this protein in the generalist necrotroph *Sclerotinia sclerotium* (SsSaxA) functions as an ITC hydrolase capable of degrading both aliphatic and aromatic ITCs in Arabidopsis (Chen et al. 2020). We therefore designated this protein BcSaxA and pursued further validation of its role in host-specific responsiveness and ITC hydrolysis in Botrytis.

### BcSaxA is an isothiocyanate hydrolase with host-specific expression

BcSaxA is located within a four-gene cluster on chromosome 6 that includes a predicted transcription factor, a Major Facilitator Superfamily (MFS) transporter (BcmfsG) previously shown to mediate ITC efflux (Vela-Corcía et al. 2019), and an additional uncharacterized ꞵ-lactamase (Figure 7A). Although BcmfsG was not detected in our proteomics dataset (likely reflecting the known underrepresentation of membrane transporters in proteomic workflows), its genomic proximity to BcSaxA, together with its established role in ITC tolerance, supports the hypothesis that this cluster functions cooperatively in detoxification of ITCs.

**Figure 7.**
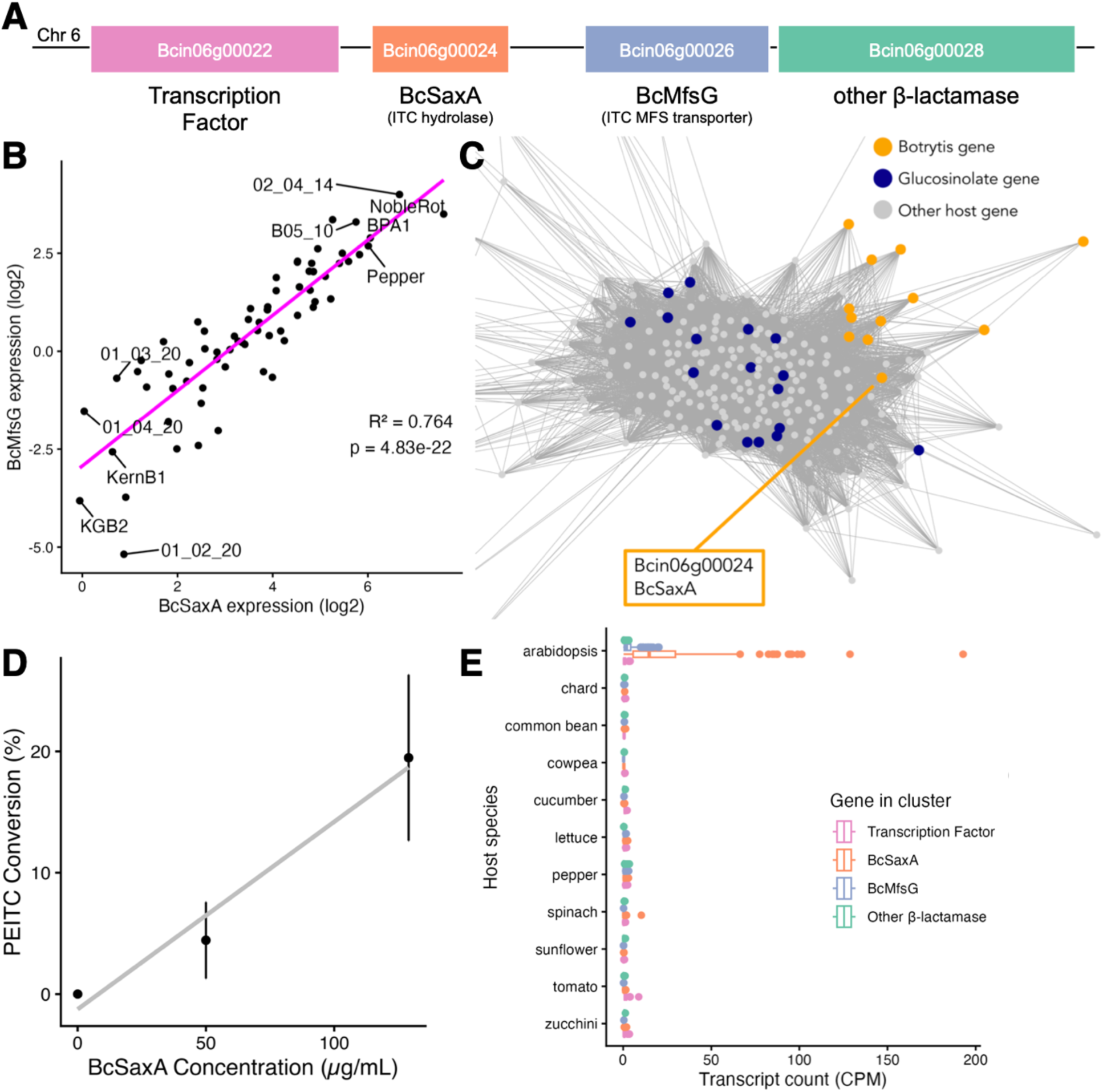
BcSaxA functions as an ITC hydrolase. A) Map of the ITC-detoxification cluster in the Botrytis genome. B) Expression of BcSaxA and BcMfsG for diverse Botrytis isolates during infection of Arabidopsis Col-0. Normalized expression values are log transformed. The 5 lowest expressing and 5 highest expressing isolates of BcSaxA are labeled with the isolate name C) Cross-kingdom gene co-expression network showing Botrytis beta-lactamase co-expresses with core glucosinolate pathway genes in Arabidopsis during infection of Col-0. Edges are drawn for PCC > 0.85 for visualization. D) *In vitro* conversion rate of phenethyl ITC by His-tag purified BcSaxA as measured by HPLC. Error bars denote standard deviation (n=4). E) Expression of Botrytis genes in the ITC-detoxification cluster from diverse isolates of Botrytis when infecting Arabidopsis and 10 additional eudicot hosts.

To assess whether expression of this gene cluster is specific to GSL-containing host plants and how it varies across diverse Botrytis isolates, we analyzed two published RNAseq datasets profiling co-transcriptomes across 72 isolates during infection of Arabidopsis (Zhang et al. 2019) and across 10 additional eudicot host species (Singh et al. 2026b). Across these datasets, expression of BcmfsG and BcSaxA was largely restricted to infection of Arabidopsis with little to no expression on all other host plants that do not contain isothiocyanates or glucosinolates (Figure 7B). In contrast, the predicted transcription factor and the additional ꞵ-lactamase Bcin06g00028 showed minimal expression under all conditions examined. Among the 72 isolates, the expression of BcSaxA and BcmfsG were positively correlated, although the magnitude of induction varied substantially between isolates (Figure 7C). This pattern suggests that BcmfsG and BcSaxA act in concert to mediate ITC tolerance in an isolate-dependent manner.

To provide more context on how Arabidopsis GSL metabolism shapes Botrytis responses, we leveraged Arabidopsis-Botrytis co-transcriptome data to identify fungal genes correlated with plant GSL biosynthetic genes expression. This analysis generated a cross-kingdom co-expression network in which several Botrytis genes, including BcBot1-5 and BcSaxA, clustered with core GSL pathway genes (Figure 7A; Supplementary Table 8). These results indicate that at the transcript level, BcSaxA is also responsive to host activation of the glucosinolate pathway.

In the proteomic data, BcSaxA was shown to accumulate when Botrytis infects Col-0 but not tgg1/2 (Figure 6), indicating responsiveness to glucosinolate breakdown products. To test whether BcSaxA directly metabolizes ITCs like the Sclerotinia homologue, we expressed and purified His-tagged BcSaxA and performed in vitro enzyme assays using phenethyl ITC as a substrate across a range of enzyme concentrations. Product formation, quantified by HPLC, demonstrated that BcSaxA directly catalyzes ITC conversion (Figure 7B).

Because the botrydial (BcBot) gene cluster also emerged from this cross-kingdom co-expression network, we examined BcBot protein abundance across infections of the Arabidopsis GSL genotypes. In contrast to BcSaxA, BcBot proteins did not show consistent changes when GSLs or their breakdown products were perturbed, suggesting that activation of this pathway is not strictly dependent on GSL-derived metabolites (Supplementary Figure 6A). This is consistent with prior reports of broad botrydial induction across hosts (Muhich et al. 2026; Singh et al. 2026b). Moreover, BcBot protein abundance did not correlate with BcSaxA levels (Supplementary Figure 6B), indicating that any relationship between these pathways is likely indirect and/or context dependent. Additional Botrytis network members showed modest genotype-dependent variation in their abundance (Supplementary Figure 7).

## Discussion

Plant-pathogen interactions involve coordinated reprogramming of pathways in both organisms at the transcript, protein, and metabolite level. Here, our time course analysis demonstrates that Arabidopsis mounts an organized proteomic response to Botrytis centered on specialized defense metabolism. Across the interaction, camalexin and indolic GSL-associated proteins were broadly induced, while many aliphatic GSL biosynthetic proteins decreased in abundance during infection (Figure 2). While the present analysis used whole leaf tissue, more work is needed to better describe potential variation of these patterns at the cellular level. Previous work has shown that camalexin is induced in response to diverse isolates while both indolic and aliphatic glucosinolates are decreased immediately in the vicinity of the developing lesion (Kliebenstein et al. 2005). This discrepancy between aliphatic and indolic GSLs with our proteomic results may be partially explained by the strong upregulation observed here in the atypical myrosinase PEN2 that primarily catabolizes indolic GSLs in cell-specific responses to pathogen attack (Bednarek et al. 2009). Overall, the host proteomic signatures described here reflect both specific activation of defense pathways and further evidence for spatially localized GSL turnover.

Using a combination of Arabidopsis defense metabolite mutants with co-proteomics identified a single protein that appears to be responsive to in planta defense metabolite variation. This host-defense metabolite dependent Botrytis protein, BcSaxA, a metallo-ꞵ-lactamase with demonstrated in vitro activity against ITCs (Figure 7D). The sax operon was first characterized in *Pseudomonas syringae* and allows for growth on Arabidopsis extracts containing ITCs (Fan et al. 2011). Interestingly, SaxA activity specifically creates an amine product that could indicate that the enzyme is both detoxifying isothiocyanates and providing a new carbon and/or nitrogen source for the pathogen (Unger et al. 2024). Subsequent analysis of seven different ITC hydrolases from various bacteria revealed similar activities on 6 structurally distinct ITCs, indicating that the enzyme functions across most ITCs (Van Den Bosch et al. 2018). Further biochemical characterization will be necessary to determine the precise catalytic activities of BcSaxA. Intriguingly, ITCs are almost entirely constrained to the Brassicales, which invokes evolutionary questions regarding the retention of specialized detoxification genes in generalist pathogens. If some mechanisms are only advantageous on certain hosts, it is unclear how selection maintains these genes across a broad host range. One possibility is that these proteins possess broader biochemical activities, while another is that they are used only under highly specific host chemical environments that are nonetheless common enough to maintain selective pressure.

A central challenge in plant-pathogen interactions is unraveling each organism’s mechanisms of sensing the other and responding by upregulating appropriate downstream pathways. Molecular signals from the plant must play a role in eliciting these responses in the pathogen. Using GSL mutants in Arabidopsis, we provide in planta evidence that the lineage-specific ITC breakdown products of GSLs are required for induction of BcSaxA. While BcSaxA was only induced on GSL-containing hosts (Figure 7E), its abundance also varied depending on host GSL composition (Figure 6). In particular, higher BcSaxA accumulation in Col-0 compared to Ler, Sha, and the AOP2 transgenic line may indicate the Botrytis mechanism inducing BcSaxA is more responsive to methylsufinyl ITCs than alkenyl ITCs. However, it is not presently known how Botrytis specifically detects and responds to isothiocyanates or other host-specific metabolites like camalexin, tomatine, resveratrol, etc. At a broad level, pathogen perception to host-specific environments and induction of tolerance mechanisms remain unresolved. Botrytis has over 1000 surface-associated proteins, many of which may contribute to host sensing and recognition (Escobar-Niño et al. 2021). Downstream signaling likely involves a combination of cAMP, MAP-kinase, histidine-kinase, and calcium-depending signaling pathways (Liñeiro et al. 2016). Future work centered on how Botrytis detects and responds to host-specific defense metabolites will be key to help better understand the precise mechanics of generalist host-pathogen interactions.

Together, this demonstrates that the Arabidopsis-Botrytis interaction is shaped by reciprocal metabolic strategies from both organisms during infection. Though GSLs are often thought of as more static phytoanticipins, GSL pathway enzymes are in fact regulated in an organized manner in response to Botrytis. Conversely, Botrytis responds to these distinct host chemical signals with its own tailored mechanisms, highlighting a level of metabolic specificity not typically associated with generalist pathogens.

## Methods

### Plant and fungal materials and growth conditions

Arabidopsis thaliana genetic material included the 3 wild type accessions Columbia (Col-0), Landsberg erecta (Ler), and Shahdara (Sha). In addition, we utilized 5 mutants with altered glucosinolate profiles in the Col-0 background: AOP2 (Li and Quiros 2003), *tgg1/2* (Barth and Jander 2006), *myb28, myb29,* and *myb28/29* (Sønderby et al. 2010). Genetic resources of Botrytis cinerea included reference isolate B05.10 and was stored at -80°C as a spore stock in 60% glycerol.

For each co-proteomics experiment, Arabidopsis seeds were cold-stratified in deionized water for 3 days at 4°C, then grown for 8 weeks in cell flats containing Sunshine Mix #1 (Sun Gro Horticulture, Agawam, MA) horticulture soil in a growth chamber at 20°C with 10h of light at 100-120uE light intensity. They were watered twice weekly with deionized water for the first 2 weeks, then with nutrient solution (0.5% N-P-K fertilizer in a 2:1:2 ratio; Grow More 4-18-38). Botrytis spores were diluted 1:10 (v/v) glycerol stock in sterile grape juice and grown on potato dextrose agar plates for 2 weeks to obtain spores for infection assays.

### Detached leaf assay and protein extraction

To reveal the Arabidopsis-Botrytis co-proteomics dynamics across early infection, we first conducted a time course of Arabidopsis leaf infection. Leaves were inoculated with Botrytis isolate B05.10 spores following a previously described infection assay (Denby et al. 2004; Corwin et al. 2016; Soltis et al. 2019). Eight week old detached leaves distributed across trays containing 1 cm of 1% phytoagar with humidity domes at room temperature. Botrytis spores were collected in water and diluted to 10 spores/µL in 50% grape juice. Four evenly dispersed inoculum droplets of 4 µL each were placed onto each leaf lamina to ensure sufficient Botrytis infection signal. Mock (blank inoculum) and Botrytis infected leaf tissue were collected at 0, 8, 16, 24, 32, 40, and 48 hours post inoculation (HPI) with 3 biological replicates for each timepoint and treatment. At each timepoint, leaves were flash frozen in liquid nitrogen and stored at -80°C until protein extraction.

To reveal the effects of glucosinolates on Arabidopsis-Botrytis co-proteomics, we analyzed infected leaf tissue of genotypes with differing GSL profiles represented by 3 wild-type accessions and 5 mutant lines of Arabidopsis (Table 1). As above, 8 week old detached leaves of the Arabidopsis genotypes were inoculated with B05.10 spores or mock, with 3 biological replicates each. At 48 HPI, leaves were flash frozen in liquid nitrogen and stored at -80°C until protein extraction.

To extract protein for all samples, 100 mg of full frozen leaf was homogenized in 500 μL buffer (5% (w/v) SDS, 50 mM triethyl ammonium bicarbonate) and incubated on ice for 2h. Extract was centrifuged at 20,000 x *g* for 15 min. Supernatant was centrifuged again for 5 min to remove any remaining cell debris. Protein concentration and purity was assessed on a nanodrop spectrophotometer. The protein extracts were frozen at -80°C until proteomic analysis.

### Defense metabolite genotype co-proteomic data collection

Defense metabolite genotype co-proteomics was performed at the UC Davis Proteomics Core Facility. Protein extracts were reduced by DTT (10 mM). Iodoacetamide (55 mM) was then added to alkylate the reduced sulfhydryl. After washing and dehydration with acetonitrile, the proteins were digested with trypsin and subjected to MS analyses.

Tandem mass spectrometry (MS/MS) was performed on an Orbitrap Fusion Lumos Tribrid mass spectrometer (Thermo Fisher Scientific) coupled to an Ultimate 3000 RSLCnano system (Thermo Fisher Scientific). Proteins were lyophilized and redissolved in 15 μL of solvent (2% CH3CN/0.1% formic acid [FA] in ddH2O). Samples (3 μL) were loaded on an Acclaim PepMap 100 precolumn (100 μm × 2 cm, C18, 5 μm, Thermo Scientific) and eluted on an Acclaim PepMap 100 analytical column (75 μm × 15 cm, C18, 3 μm, Thermo Scientific) with a 60-min gradient. FA (0.1%) and CH3CN/H2O (4:1, v/v) with 0.1% FA were used as mobile phase A and B, respectively. The loading buffer was the same as the mobile phase A. The gradient was: 5 min at 4% B, 31-min linear gradient from 4% B to 30% B, 7-min linear gradient from 30% B to 50% B, 3-min linear gradient from 50% B to 99% B. After washing the column for 4 min with 99% B, the buffer was decreased to 4% B in 1 min and the column was reconditioned with initial gradient for 9 min. The column temperature was 35°C. The flow rates through the trap column and analytical column were 3 μL/min and 0.3 μl/min, respectively. The MS/MS experiment was operated in the data-dependent mode with automatic switching between MS and MS/MS scans with a cycle time of 3 s. The full MS scans were operated over a mass range of *m*/*z* 350–1800 with detection in the Orbitrap (120K resolution). Fragmentation of ions was achieved with high-energy collisional dissociation (30% collision energy) and the spectra were acquired in the Orbitrap (30K resolution) with a maximum injection time of 100 ms.

The LCMS raw files were processed with Proteome Discoverer 2.5 (Thermo Fisher) using the integrated SEQUEST engine. All data was searched against Uniprot reference proteome, UP000006548 (*Arabidopsis thaliana* Columbia) and UP000001798 (*Botrytis cinerea* B05.10). Peptide tolerance and fragment mass tolerance were set to 10 ppm. Trypsin was specified as enzyme, cleaving after all lysine and arginine residues and allowing up to two missed cleavages. Only variable modifications were used and they included the following: carbamidomethylation of cysteine, protein N-terminal acetylation, oxidation of methionine. The false discovery rate (FDR) was set to 1% on peptide spectrum match (PSM), PTM site and protein level.

### Protein extraction and sample preparation for time course proteomics

One hundred microliters of pre-warmed (65°C) extraction buffer (4% (w/v) SDS; 30% (w/v) sucrose; 50mM Tris-HCl, pH = 8.0; 1x phosphatase inhibitor cocktail) was added to ∼100 mg of finely-ground tissue and briefly vortexed until completely mixed. Sample was then incubated for 10 minutes at 65°C and cooled down for 5 minutes at room temperature. 100µL of buffer-saturated phenol (pH = 8.0, Thermo Scientific) was added to the protein extract and sample was vortexed for 1-2 minutes. Protein extract was centrifugated at 10,000xg for 5 minutes and organic (upper) layer was transferred to a new tube. A second round of phenol extraction was performed on the remaining protein extract, and the organic phase was transferred and pooled with that of the first phenol extraction. Extracted proteins were precipitated from organic phase at -80°C for 2 hours in 1mL of 0.1M ammonium acetate in ice-cold methanol. Samples were centrifuged for 5 minutes at 10,000xg, 4°C and supernatant was discarded. Pellet was washed twice in ice-cold 100% acetone (i.e., 1mL of acetone was added and pellet was resuspended by probe sonication, followed by a 5-minute incubation at -20°C and centrifugation for 5 minutes at 10,000xg, 4°C). Finally, pellet was washed once on ice-cold 70% methanol and washed pellet was set to air dry for ∼10 minutes with care not to over dry.

Extracted proteins were further cleaned using the Protein Aggregation Capture (PAC) method. Specifically, samples were probe-sonicated in 4% SDS; 50mM Tris-HCl, pH = 8.0, total protein was quantified using BCA (Thermo Scientific). Three hundred micrograms of sample was mixed with MagReSyn Hydroxyl beads (Resyn Biosciences) at a 1:2 (sample:bead) ratio. Acetonitrile (ACN) was added until the ACN final concentration was 70%. Two PAC rounds, consisting of 1 minute with gentle agitation (∼1000rpm) followed by 10 minutes of incubation at room temperature with no shaking were carried out. Beads were washed once with 100% ACN, and twice with 70% methanol. Proteins were digested into peptides in 50mM TEAB on-beads at 37°C overnight using 3µg of Trypsin (Roche). A second digestion round was performed the next day using 500ng of Trypsin and 50ng of Lys-C (Wako). Peptides were eluted from beads twice using 100µL of 1% Trifluoroacetic acid (TFA), then vacuum-dried until almost dry.

### LC-MS/MS for time course proteomics

Chromatography was performed on a Thermo Vanquish Neo UHPLC in “heated trap-and-elute, backward flush” mode. Peptides were desalted and concentrated on a PepMap Neo trap column (300 µM i.d. x 5 mm, 5 µm C18, 100 Å µ-Precolumn, Thermo Scientific) at a flow rate of 5 µL min^-1^. Sample separation was performed on a 60 cm Aurora Frontier Column (Ionopticks) with a flow rate of ∼300 nL min^-1^ over a 180 min reverse phase active gradient (80% ACN in 0.1% FA, from 7.2% to 24% over 109 min and from 24% to 44% in 11 min). Followed by a column/trap wash at 78% ACN for 10 min. Eluted peptides were analyzed using a Thermo Scientific Orbitrap Exploris 480 mass spectrometer with a FAIMS pro Duo interface installed, which was directly coupled to the UHPLC through an Easy Spray Ion source (Thermo Scientific). Data dependent acquisition was obtained using Xcalibur 4.0 software in positive ion mode with a spray voltage of 1.8 kV, a capillary temperature of 280 °C, an RF of 45, FAIMS compensation voltage of -45, and a total carrier gas flow of 4.2 l/min. MS1 spectra were measured at a resolution of 60,000, an automatic gain control (AGC) of 2.25e6 with maximum injection time set on “Auto”, and a mass range of 400-900 m/z. Forty two DIA acquisition windows of 12m/z with 1Da overlap were measured at a resolution of 30,000, an automatic gain control (AGC) of 8e6 with auto maximum ion time, over a precursor mass range of 400-900 m/z. A scan range 145-1450 m/z and normalized collision energy of 29 were used.

Spectra were analyzed using Spectronaut version 19 (Biognosys). Spectra were searched, using the Pulsar search engine against the Arabidopsis thaliana ARAPORT11 reference annotation from The Arabidopsis Information Resource (TAIR, www.arabidopsis.org), and the Botrytis cinerea B05.10v2 annotation (NCBI accession GCA000143535.2). The spectra search was performed on “direct DIA” mode. Carbamidomethyl cysteine was set as a fixed modification while methionine oxidation and protein N-terminal acetylation were set as variable modifications. Digestion parameters were set to “specific” and “Trypsin (Full)”. Up to two missed cleavages were allowed. A false discovery rate, calculated in Percolator using a target-decoy strategy, of less than 0.01 at both the peptide spectral match and protein identification level was required.

### Proteomic data analysis

Raw proteomic time-course data recovered 11,712 Arabidopsis proteins and 1,548 Botrytis proteins. To reduce the influence of stochastic peptide detection, proteins quantified in only a single replicate were considered unreliable and their values were recoded as zero. Low abundance proteins were further filtered by retaining only those detected at least two replicates within any treatment x time combination. After filtering, 11,669 Arabidopsis and 1,537 Botrytis proteins remained.

All protein models were conducted with the R package glmmTMB using a negative binomial distribution (Brooks et al. 2017). For each model, the resulting p-values were adjusted for multiple comparisons using the false discovery rate (FDR) correction (Benjamini and Yekutieli 2001). For Arabidopsis time-course samples, individual proteins were modeled using the formula:

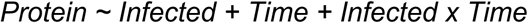

After removing 27 outlier proteins, hierarchical clustering was performed on the remaining proteins significant (p < 0.05) for the ‘infected’ or ’infected x time’ term using complete linkage. For visualization, extreme log2FC values were capped at ±5. Clusters were identified using the dynamicTreeCut R package (Langfelder et al. 2008) with a minimum cluster size of 30. GO enrichment (biological process ontology) for each cluster was performed against all detected Arabidopsis proteins using the topGO R package (Adrian Alexa 2017). Significant GO terms (p < 0.05) were filtered for redundancy both manually and using REVIGO (Supek et al. 2011).

To assess for Botrytis protein dynamics in the time course samples, individual proteins were modeled at each timepoint using the formula:

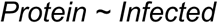

First detection time was defined for each protein as the earliest timepoint at which p < 0.05 for the mock versus infected comparison, provided the protein remained detected at the subsequent 8- hour timepoint to reduce noise. Hierarchical clustering of all detected Botrytis proteins was performed using average linkage and Euclidean distance based on protein abundances.

For modeling Arabidopsis protein shifts across Arabidopsis defense metabolite genotypes during infection, genotypes were split into 4 groups (accessions, Col-0 vs *tgg1/2*, Col-0 vs AOP2, and Col-0 vs *myb28, myb29, and myb28/29*). Each protein was modeled within each group with the formula:

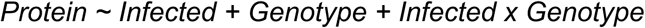

For modeling Botrytis protein shifts on the Arabidopsis genotypes, the genotypes were split into the same 4 groups as above. Proteome data for infected samples was modeled within each group with the formula:

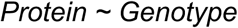

### Cross-kingdom co-expression network analysis

Normalized gene expression data from previously published Arabidopsis-Botrytis co-transcriptomes were utilized to generate cross-kingdom co-expression networks (NCBI SRA PRJNA473829, Zhang et al. 2017). This included transcripts of both Arabidopsis Col-0 and 96 isolates of Botrytis cinerea at 16 HPI. Co-expression networks were constructed as described in Wisecaver et al. 2017. In brief, Pearson correlation coefficients were calculated for all host and Botrytis gene expression values across the isolates. Correlations were ranked and mutual rank was calculated for each gene pair. Mutual ranks were converted to network edge weights using an exponential decay rate of 50. Overlapping modules of tightly co-expressed genes were called using ClusterONE (Nepusz et al. 2012). Resulting networks were screened for those containing both Arabidopsis glucosinolate genes and any Botrytis genes.

### Protein Expression and Purification

The open reading frame of BcSaxA (Bcin06g00024) was amplified from B. cinerea B05.10 cDNA and cloned into the expression vector pET-28a(+) (Twist Biosciences). The recombinant plasmid was expressed in *E. coli* strain BL 21 (DE 3). The cells were incubated for 2 hours on a shaker at 37°C and 220 rpm, until an OD600 of 0.35-0.45 was reached, and subsequently induced by adding isopropyl β-D-1-thiogalactopyranoside to a final concentration of 0.5 mM and set on a shaker at 16-18°C 200 rpm for 24 h. *E. coli* cells were harvested by centrifugation at 8000 x *g* at 4°C for 15 mins. The pellets were washed, resuspended in Lysis solution (50 mM H2NaPO4, pH 7, 200 mM NaCl, 10% glycerol, 15 mM Imidazole, 1 mM DTT, 0.5 mM PMSF), sonicated on ice, and centrifuged at 16,000xg for 15 min at 4°C. The crude protein was purified with Ni-NTA resin for 6xHis tagged proteins (Qiagen, Hilden, Germany) following the protocol described by the manufacturer. An SDS-PAGE gel was run to verify the size and purity of the protein, and a Bradford assay was used to determine the concentration of the purified protein.

### Enzyme assay and HPLC

Enzyme assays of the purified BcSaxA protein were performed with 2-phenethyl isothiocyanate (2PE-ITC) in DMSO. The reaction was conducted at 25°C with the following conditions: 490 mM 2PE-ITC, 40 mM KP buffer (KH2PO4-K2HPO4, pH 7.0), 10 µM ZnSO4, 10 µL purified recombinant protein at three concentrations (0, 50, 129 µg/mL) and a heat inactivated control. The final volume of the reaction mixture was diluted to 100 µL with water. The reaction proceeded for 15 minutes, then was terminated with the addition of 100 µl methanol. Products were centrifuged for 20 minutes at 20,000 x *g* and 4°C.

The products of the ITCase enzyme assay with 2PE-ITC were analyzed using a 1100 series HPLC system (Agilent Technologies) with a diode array detector. Separation of ITCs was achieved on a LiChrospher 100 RP-18 column (250 x 4.0 mm, 5 µm, Agilent Technologies) with water (A) and acetonitrile (B) at a flow rate of 1.0 ml min⁻¹ using a gradient as follows: 0-4.99 min, 5% B; 5-17.99 min, 15% B in A; 18-22.99 min, 95% B in A; 23-25 min, 5% B. The peak area of the 2PE-ITC signal at 229 nm and 245 nm was identified at 17.9 min and the breakdown product peak was identified at 17.5 min. 2PE-ITC was purchased from Sigma Aldrich (Steinheim, Germany).

## Data Availability

Raw proteomic data will be made available at ProteomeXchange upon publication.

## Acknowledgements

This work was supported by the NSF award IOS 2020754 for DJK. The authors acknowledge HPC@UCD for providing computational resources that have contributed to the research results reported in this paper.

## Supplementary Figures

**Supplementary Figure 1.**
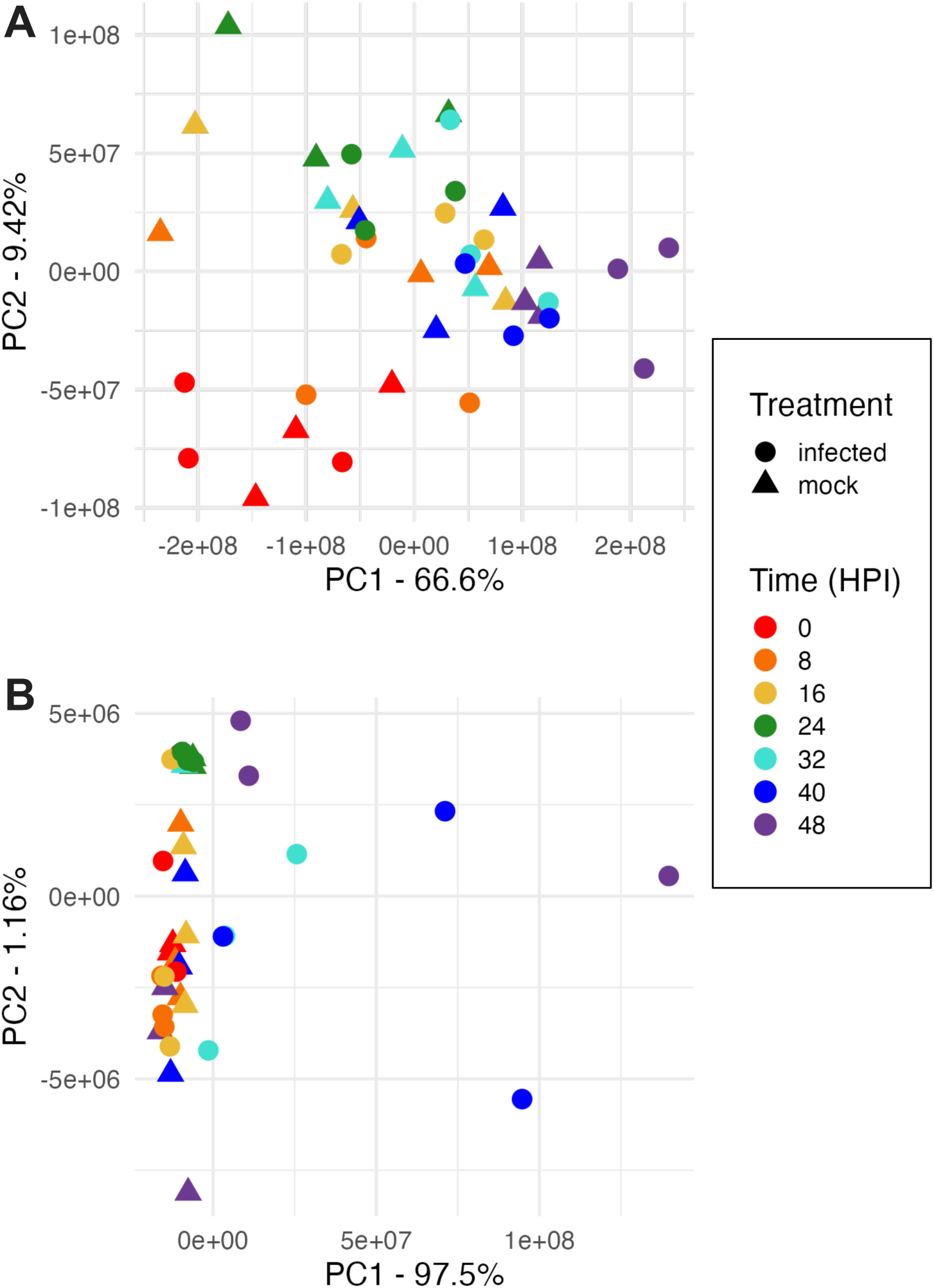
Principal component analysis (PCA) of A) Arabidopsis and B) Botrytis proteomes across the 48H infection time course. Circles represent infected samples and triangles reflect mock control samples. Time (HPI, hours post inoculation) is shown by color.

**Supplementary Figure 2.**
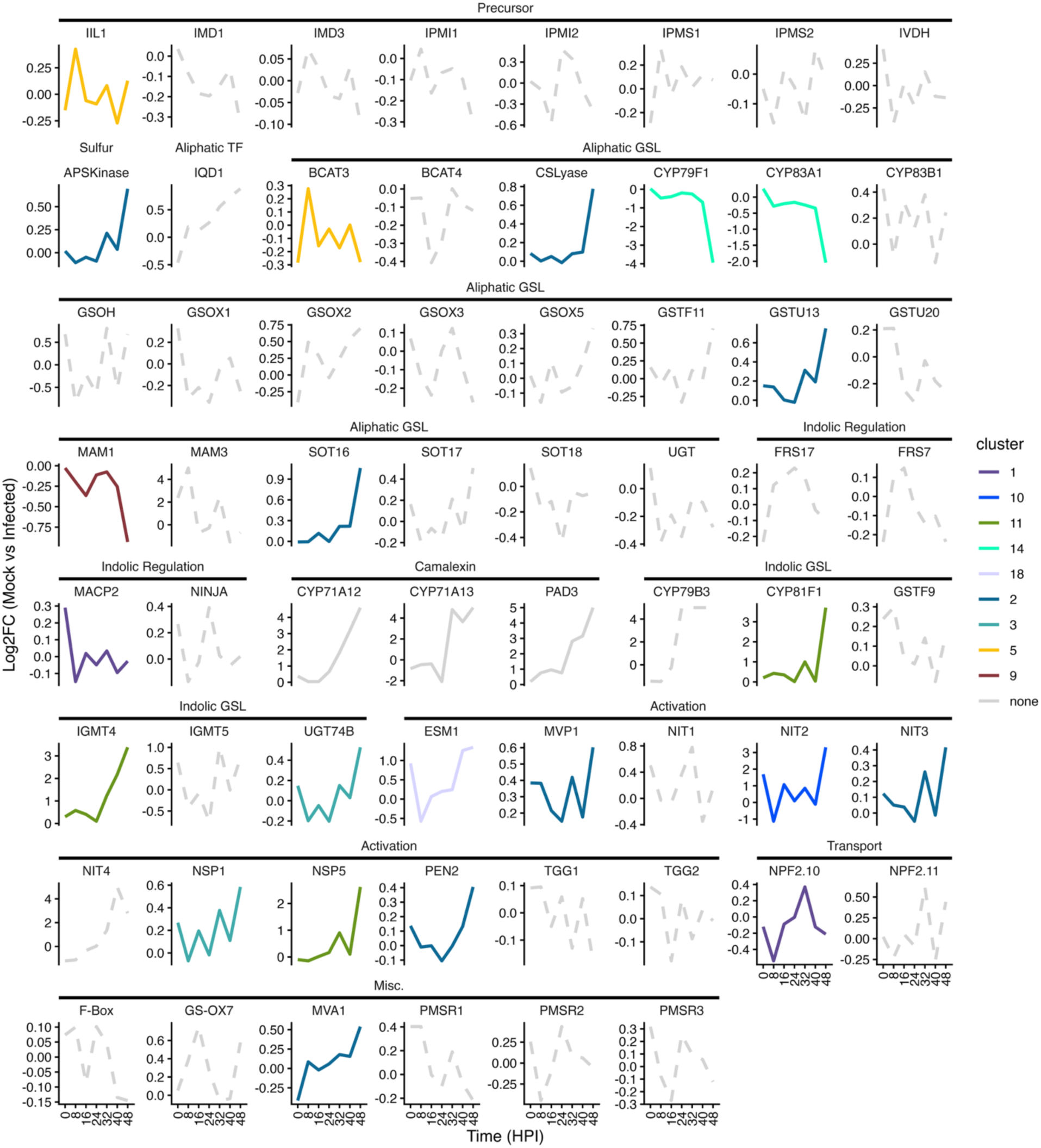
Arabidopsis glucosinolate and camalexin pathway protein abundances across time course infection with Botrytis isolate B05.10. Proteins are grouped by pathway membership. Lines are colored by cluster memberships defined by HCA outlined in Figures 1 and 2, where lines in grey were excluded from HCA. Solid lines indicate the protein level responded significantly (p < 0.05) to infection or infection x time. The following relevant pathway proteins are not shown due to low abundance: APK3-4, SDI2, MYB28, MYB29, MYB76, MYB115, MYB118, AOP2-3, BZO1, CYP79F2, GSOX4, MYB34, MYB51, MYB122, OBP2, CYP71A27-28, CYP81F2-4, CYP79B2, IGMT1-3, OMT, ESP, NSP2-4, TGG4-5.

**Supplementary Figure 3.**
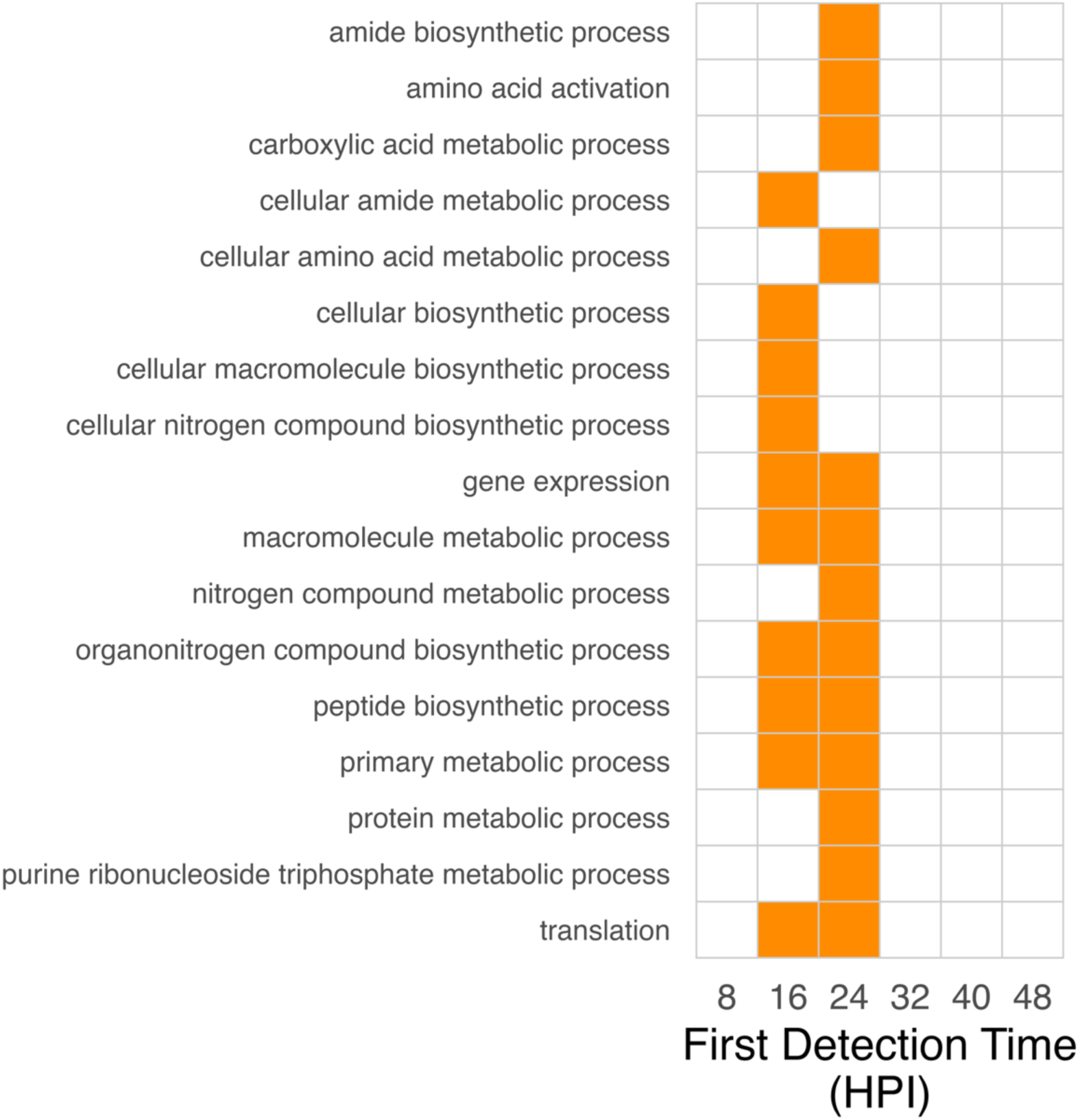
GO enrichment analysis of proteins grouped by their first detection time. Significantly enriched (p < 0.01) GO terms are shown in orange. GO enrichment was conducted with the BP ontology and redundant terms filtered. GO enrichment was performed against all detected Botrytis proteins as a background.

**Supplementary Figure 4.**
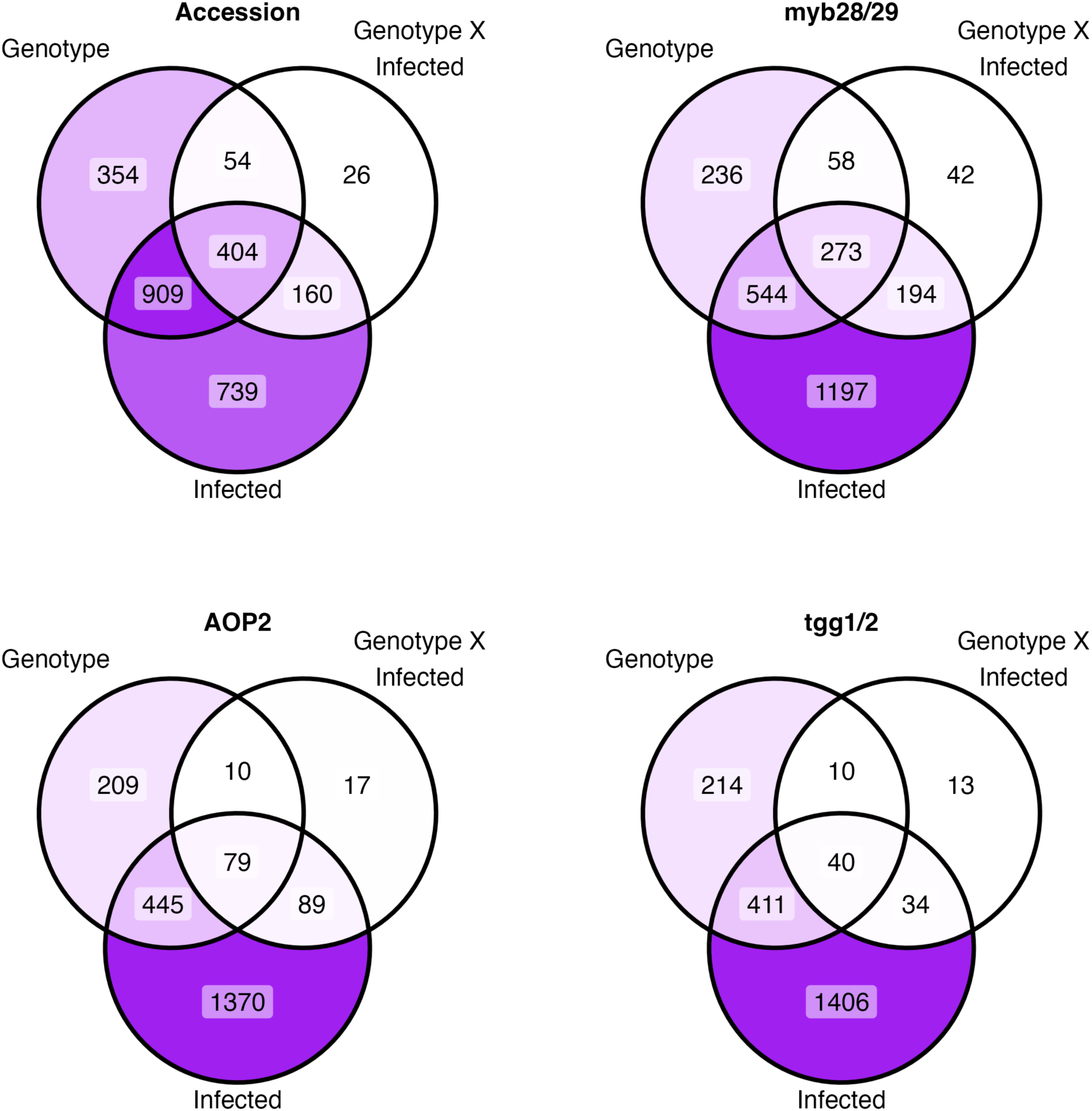
Count of Arabidopsis proteins with significantly different accumulation (p < 0.05) across different Arabidopsis genotype comparisons during infection with Botrytis. “Accession” compares Col-0, Ler, and Sha; “myb28/29” compares Col-0, *myb28, myb29* and *myb28/29*; “AOP2” compares Col-0 and AOP2; “tgg1/2” compares Col-0 and *tgg1/2*.

**Supplementary Figure 5.**
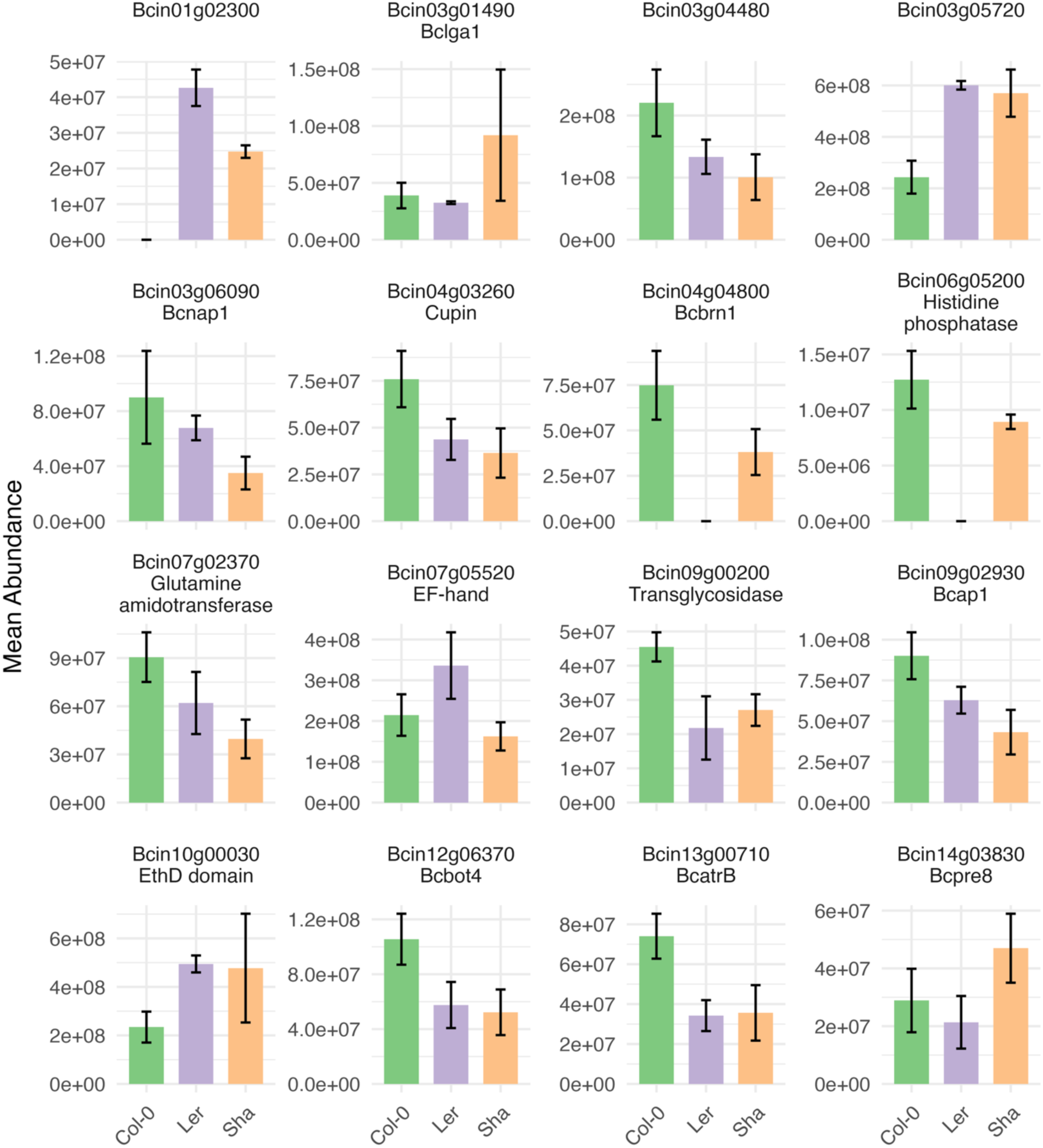
The 16 Botrytis proteins significantly altered during infection of multiple WT Arabidopsis accessions (p < 0.05, >= 2FC). Mean and SD of protein abundances for each significant comparison are shown.

**Supplementary Figure 6.**
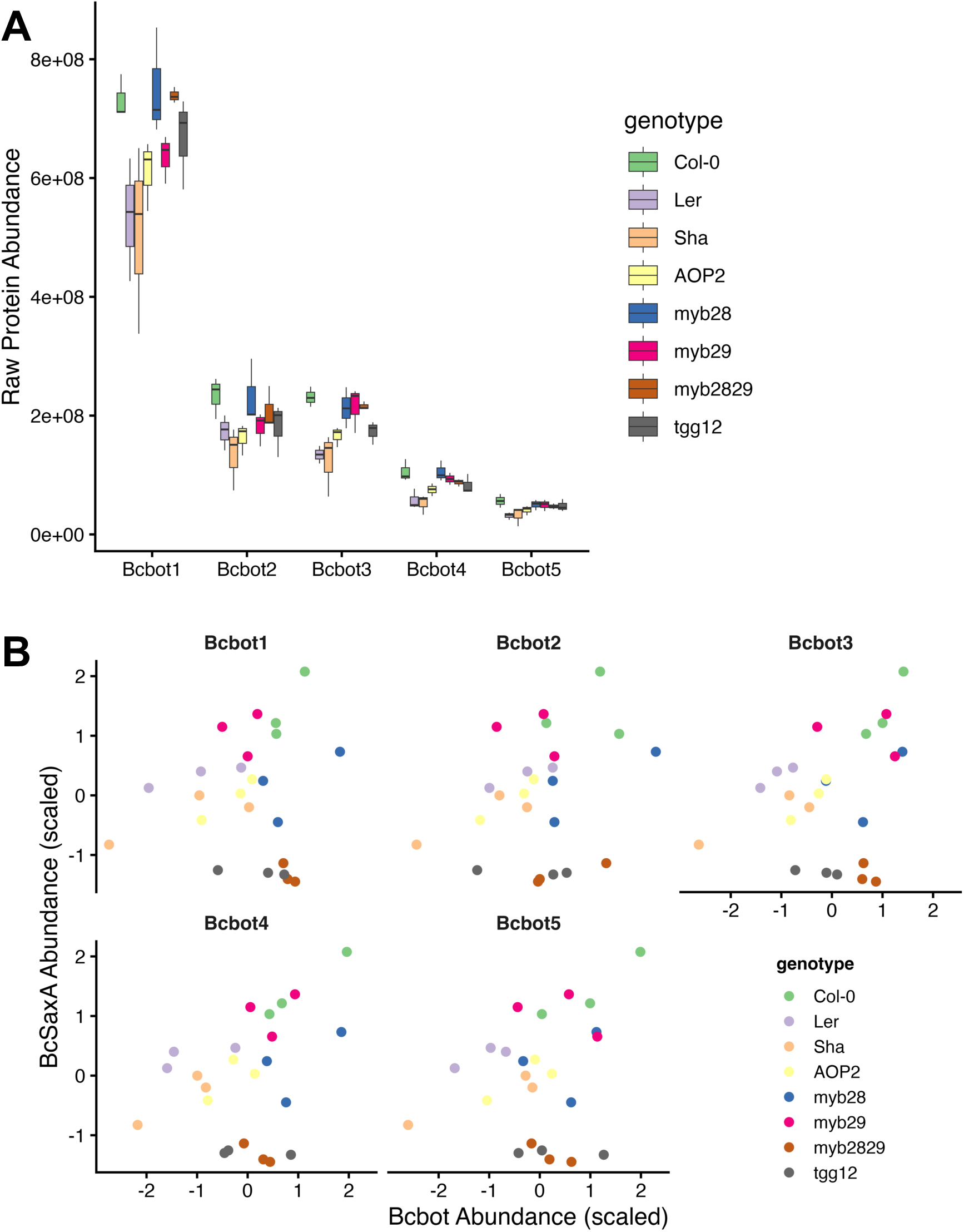
A) Bcbot raw protein abundance during infection of Arabidopsis genotypes. All 5 proteins in the Bcbot cluster are shown. B) Bcbot vs BcSaxA protein abundance correlations during infection of Arabidopsis accessions and GSL mutant lines. Abundance values are scaled within each protein.

**Supplementary Figure 7.**
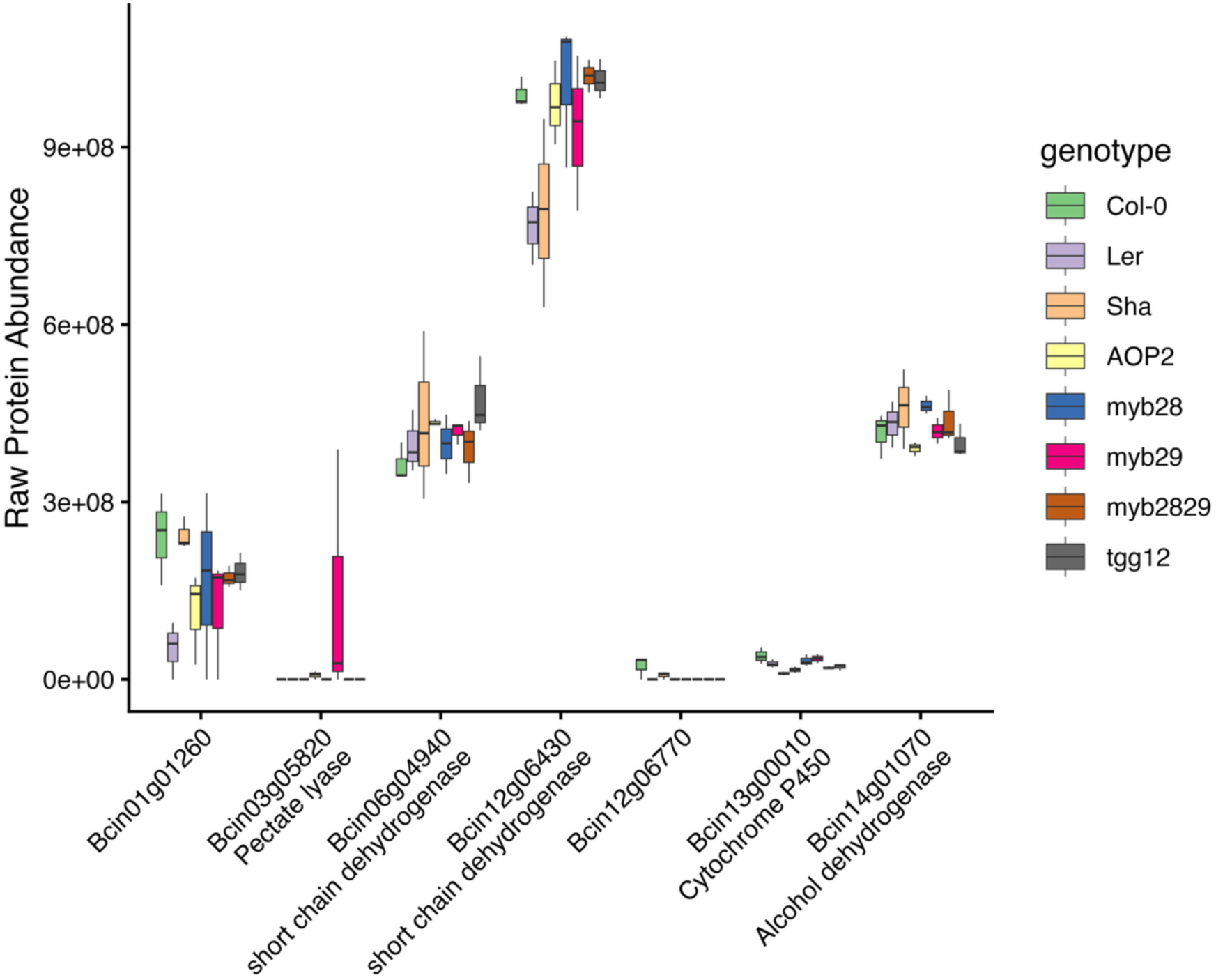
Protein abundances of other Botrytis network members recovered in the cross-kingdom gene co-expression network (Figure 7A). These Botrytis genes were found to co-express with Arabidopsis glucosinolate genes.

